# TRPM7–Annexin A1 Mechanosensitive Pathway Drives Capillary Infiltration by Circulating Tumor Cells

**DOI:** 10.1101/2025.08.01.668151

**Authors:** Lanfeng Liang, Zhifeng Zhao, Jonathan Wei Bao Chua, Yun Chen, Xiao Song, Lina Hsiu Kim Lim, Jiamin Wu, Qionghai Dai, Andrew William Holle, Chwee Teck Lim

## Abstract

Successful metastatic dissemination requires tumor cells to overcome significant physical barriers. When circulating tumor cells (CTCs) become lodged in capillary beds, studies reveal that only a small subset can adapt to fluid shear stress (FSS) within the constricted vasculature. These rare, mechanically resilient cells may subsequently extravasate and form metastatic lesions, but details on this adaptation of the physical environment and its molecular basis are not well understood. Utilizing a microfluidics platform that mimics microcirculatory dynamics, we discover that only breast cancer cells with high metastatic potential maintain directional migration under physiologically relevant FSS conditions. This highlights the mechanical selection process of metastatic precursors during hematogenous dissemination. We identify the TRPM7-Annexin A1-actin signaling pathway as essential for overcoming physical barriers and regulating cell motility under capillary FSS. FSS prompts cytoskeletal reorganization and actin disassembly, restricting cell movement. Cancer cells respond to this by increasing mechanical loading, activating TRPM7 and triggering calcium influx, which then activates Annexin A1. This calcium-dependent protein, Annexin A1, interacts with the actin cortex to prevent FSS-induced actin disassembly, thus aiding migration. Experiments conducted in mouse liver capillaries validate the critical role of this pathway in cancer cell motility. Additionally, we propose a mechano-pharmacological strategy using FTY720 to target the TRPM7 pathway, highlighting its therapeutic potential to modify CTC receptor specificity and inhibit distant metastasis.

## Introduction

Metastasis is responsible for over 90% of cancer-related mortality^1^. For metastasis to occur, disseminating cancer cells should intravasate into the circulatory system^2^, where CTCs are dispersed by blood flow, significantly enhancing their metastatic potential. To systemically investigate the metastatic potential of human tumor cell lines, numerous studies have utilized intracardial injection to place cells directly into the bloodstream^3^ in order to mimic the behavior of CTCs during metastasis. However, not all primary tumors lead to metastatic dissemination^4^. In fact, less than 0.1% of these CTCs successfully exit the bloodstream and become potential sources of metastases^5^. Cancer cell lines capable of successfully seeding metastases are defined as highly metastatic. In contrast, less metastatic cancer cell lines fail to form tumors via intracardial injection but can successfully form tumors following subcutaneous injection^6,7^, which bypasses the circulatory system. This suggests that the circulatory system imposes significant constraints on metastatic seeding^5,8^, but the factors determining success or failure during circulation remain unclear.

One potential modulator of this phenomena is the physical microenvironment in the circulatory system. Recent advances in intravital imaging techniques have revealed that arrested CTCs can be rupture under physiological shear forces within a few hours in the microvasculature^9,10^. The FSS in capillary vessels can be as high as 5-40 N/cm^2^ compared to approximately 0.01 N/cm^2^ in primary tumor lesions and 0.02-0.5 N/cm^2^ in lymphatic vessels^11^. Previous studies suggest that the cell nucleus can expand and rupture under high FSS (20 N/cm^2^)^12^. To avoid this rupture, arrested cancer cells must migrate and escape microcirculation, enduring physical stresses from blood flow and capillary confinement^9,13,14^. It is well-established that physical environments regulate cell motility, and the adaptation of these environmental stresses might directly determine the success or failure of metastasis^15,16^. In this case, it is still unclear whether highly metastatic breast cancer cells can withstand these mechanical stresses more effectively than low- or non-metastatic cells, and whether highly metastatic breast cancer cells are equipped with special mechanisms to respond to these environmental stresses and tailor their motility. If so, we sought to determine the underlying mechanism and to see whether it could be a target to selectively eliminate highly metastatic breast cancer cells during circulating metastasis.

## Results

### Highly metastatic breast cancer cells can maintain migration under capillary bed-mimicking physical stress

While *in vivo* observations have shown that cancer cells can actively migrate in capillary vessels^9,10,17^, the molecular mechanisms giving them this ability are poorly understood, likely due to the inherent challenges in working with *in vivo* animal models. To address this, we built a microfluidic system to expose cancer cells to physiological levels of confinement and shear stresses and mimic the metastasis on microcirculation. To quantify fluid flow in capillary bed *in vivo*, we used two-photon synthetic aperture microscopy^18^ to visualize the 3D structure of mouse liver capillaries *in situ* (***Extended Data Fig. 1a***) and dynamically track blood flow (***Supplementary Video 1***). Using these capillary diameter and blood flow rate quantifications (***Extended Data Fig. 1b*** and 1c), we designed a ‘capillary bed-on-a-chip’ that recapitulated the *in vivo* capillary microenvironment (***Extended Data Fig. 1e***). This polydimethylsiloxane (PDMS)-based system used a syringe pump to generate a continuous and controllable laminar flow tuned to match the flow profile observed *in vivo*, and enabled high-resolution time-lapse microscopy of cell migration (***Extended Data Fig. 1f-k, 2a, and Supplementary Video 2***). To investigate the broad effects of capillary vessel-like confinement and shear stress on cancer cell migration, we used a panel of human breast cancer cells with different levels of metastatic potential^3^. While previous studies have found that low levels of shear stress (0.05 N/cm^2^) can promote cancer cell motility^19^, we found the opposite phenomena when cells are presented with both confinement and higher fluid shear stress (5.9∼11.8 N/cm^2^). To better quantify this reaction, we classified cells as either migratory or non-migratory based on whether they migrated over 50 µm within 8 hours (***Fig. 1a***). When comparing cells across a gradient of metastatic potential, non-metastatic cells like MCF10A were still able to form membrane protrusions (***Supplementary Video 2***), but a large proportion of these cells were unable to migrate under conditions of confinement and flow (***Fig. 1b***), and in cells that were able to migrate, their speed decreased by over 60% (***Fig. 1c-e***). On the other hand, highly metastatic cancer cells like MDA-MB-231 remained migratory under flow as they did under static conditions (***Fig. 1b***), and motile cells only displayed a reduction in migration speed of under 20% (***Fig. 1c-e***), although most of these cell lines had similar migration speeds in standard 2D culture (***Fig. 1c***). Beyond migration speed, the directionality of migration with respect to flow was also calculated (***Fig. 1f***). As expected, no cells displayed a directionality preference under static conditions (***Fig. 1g***). Interestingly, under fluid flow, cells of low metastatic potential showed preferential migration in the direction of flow while highly metastatic cells remained indifferent to flow direction (***Fig. 1h***). To determine if this specific migration is actually a function of metastatic potential, as opposed to an innate quality of specific cell lines, we used the mouse 4T1 metastatic cell line and compared it to its non-metastatic isogenic subpopulation cell lines 67NR and 168FARN. When comparing these three isogenic lines, only the metastatic 4T1 subpopulation was capable of sustained cell migration under confinement and flow (***Extended Data Fig. 2b-d***). Additionally, cells have the ability to use the confined channel and facilitate their migration via coupling retrograde actin flow^20^. Overall, when examining how cells reduce their speed in response to flow, a general trend was observed – cells with higher metastatic potential are less sensitive to the effects of fluid flow, and we termed this metastatic-potential associated migration as ‘flow-insensitive confined migration’.

**Figure 1.**
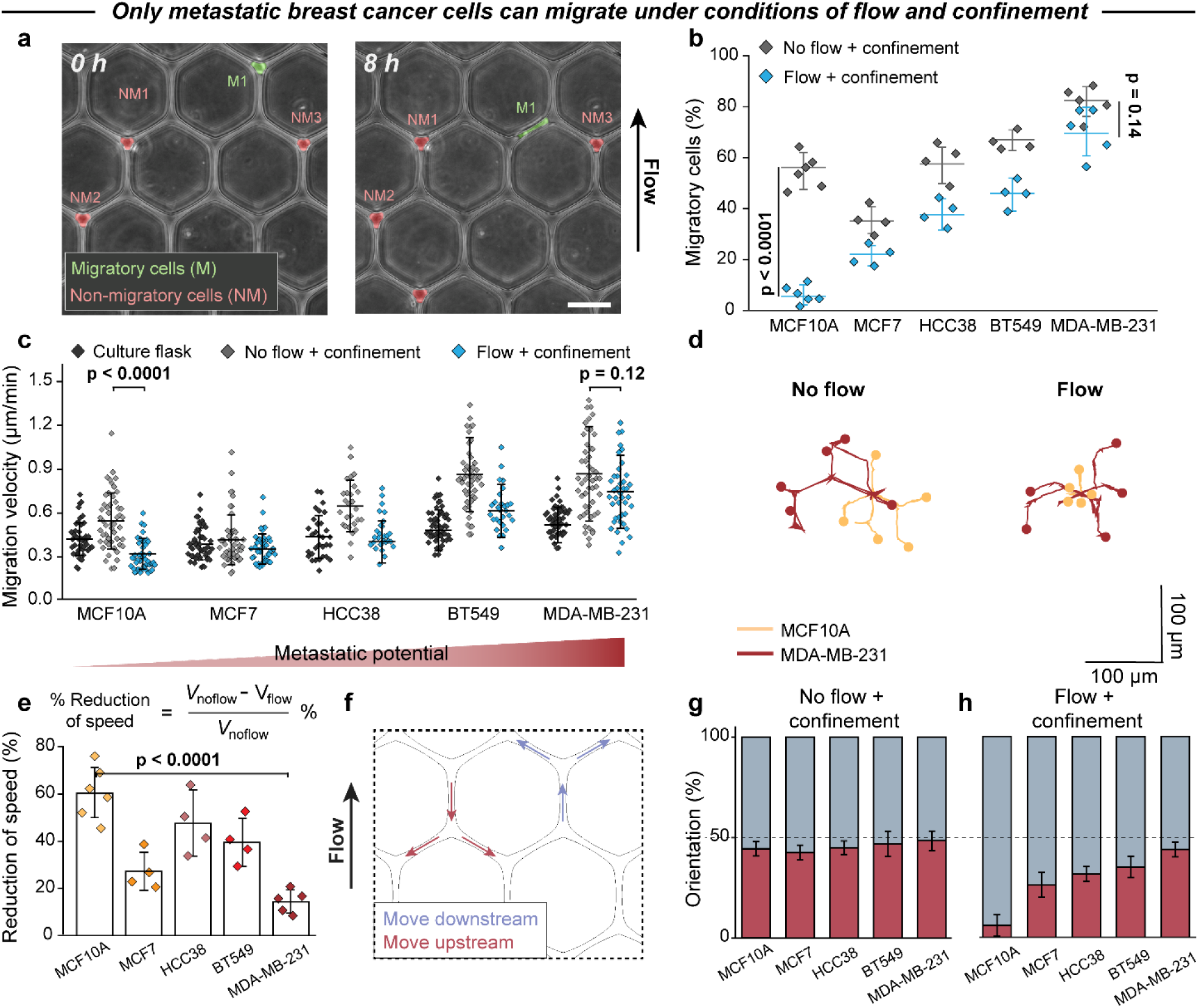
Breast cancer cells with highly metastatic potential exhibit flow-insensitive migration in confined. **a**, Definition of migratory cells: cells which migrate over than 50 μm within 8 hours. Scale bars, 50 μm. **b**, Percentage of migratory cells in breast cancer cell lines. n ≥ 3 independent experiments. **c**, Cell migration velocity under culture flask, as well as no flow and flow conditions in a confined microfluidic chip. n ≥ 30 from at least 3 independent chip experiments. **d**, Cell trajectories on microfluidic chips after 8 h. Scale bars, 100 μm. **e**, Reduction in migration speed under flow condition. **f - h**, (**f**) Orientation preference of cell migration. Gray arrows indicate ‘move downstream’, while red arrows indicate ‘move upstream’. Migration orientation preference of breast cancer cells, expressed as a percentage under no flow (**g**) and flow (**h**) condition. Data are represented as the mean ± standard deviation (n ≥ 3 independent chip experiments). Statistical analysis was performed using Kruskal-Wallis tests followed by Dunn’s test (**b, and e**) and one-way ANOVA followed by Tukey’s test (**c**).

For figuring out the impact of microchannel structure, we next designed four alternate microfluidic devices, with two allowing cells to be unconfined (***Extended Data Fig. 3a-h***), and two presenting diverse channel morphologies (***Extended Data Fig. 3i-p***). The largest reductions in migration speed in response to flow were found in the devices presenting the channel confinement, confirming that this flow-insensitive confined migration is a widely applicable phenomenon. This was not a function of cell line-specific downregulation of focal adhesions, as paxillin was found to be enriched in the leading edge of migratory cells (***Extended Data Fig. 4a-c***), suggesting that cells are actively migrating in this environment, not passively riding along with the flow. Oppositely, cancer cells can’t form paxillin-based focal adhesion if pre-coating the chip with non-adhesion agent pluronic F-127 (***Extended Data Fig. 4d***). Furthermore, to determine if extracellular matrix (ECM) composition plays a role in this process, we tested different ECM coatings inside the microfluidic channels. Both MDA-MB-231 cells and MCF10A cells didn’t alter their migratory behaviors under different ECM coating condition (***Extended Data Fig. 4e-h***).

### Cell nucleus properties do not contribute to cancer cell migration under flow and confinement

Given that confinement migration requires a dynamic reorganization of both the cytoskeleton and the nucleus, we made 3D measurements of breast cancer cells in our system and found them to exhibit reductions in both cell volume and nucleus volume under microfluidic channel (***Extended Data Fig. 5a-d***). As previous works have demonstrated the crucial role of the nucleus in responding to environmental constraint^15,21,22^, we investigated the influence of nuclear size on cell migration. Since nuclear volume has been shown to correlate with cell volume in the literature^23^, we used flow cytometry to sort cells based on size, generating three subpopulations with statistically significant differences in nuclear volume (***Extended Data Fig. 5e***). Interestingly, speed reduction as a function of flow was not influenced by nuclear volume (***Extended Data Fig. 5f***). We also actively modulated physical feature of nucleus by increasing chromatin condensation using JIB-04 or decreasing it with RGFP-966^24,25^, and we again observed no significant difference in the extent to which flow reduced migration speed (***Extended Data Fig. 5g***). We thus concluded that nuclear feature does not drive the migratory difference between metastatic and non-metastatic cells.

### Metastatic cancer cells utilize a calcium-based TRPM7 mechanism to tailor their migration under flow and confinement

Calcium plays a central role in cell migration and cancer metastasis and has proven to be a key mechanism through which cells decode information about their physical microenvironment^26,27^. Thus, we next examined the role of calcium signaling (***Fig. 2* *and Extended Data Fig. 6)***. First, to exam whether calcium is required for this migratory process, we create a calcium-free experimental environment via introducing MDA-MB-231 into either calcium-free media and media containing the calcium chelator EGTA in microfluidic chip^28^. We found that, in contrast to control media or media with physiological calcium levels, MDA-MB-231 cells almost stop their migration under calcium-free environment (***Fig. 2c-d***). Subsequently, we visualized intracellular calcium levels within the cells and observed a significant rise in response to fluid flow (***Fig. 2a-b* *and Extended Data Fig. 6a-b***). This intracellular calcium increase does not equilibrate rapidly, and the calcium flickers occur typically after one-hour post-flow induction. These calcium flickers exhibit higher activity at the leading lamella of migrating cells^27^. However, such calcium flickers were less observed on MCF10A cells (***Extended Data Fig. 6c***). Following the inhibition of calcium influx using the potent blocker GdCl_3_, MDA-MB-231 cells were no longer able to regulate their intracellular calcium levels **(*Fig. 2a-b*)**. This disruption was accompanied by a significant decrease in cell migration under flow **(*Fig. 2e-f*)**. In contrast, no such strong effects were observed when sodium or potassium ion channels were inhibited **(*Supplementary Video 3*)**, despite these channels have been previously established roles in cell migration^29^. As calcium-associated signaling networks are highly complex, with key players including intracellular calcium levels, endoplasmic reticulum (ER)-associated calcium release and storage, and stretch-activated calcium ion channels^22,30^, we performed a small pharmacological inhibitor screening on each potential mechanism. While each calcium inhibitor on its own did not affect MDA-MB-231 confinement migration, once flow was applied, the largest reductions in both the percentage of migratory cells and migration speed were observed when inhibitors targeting stretch-activated calcium ion channels were used (***Extended Data Fig. 6d-e***).

**Figure 2.**
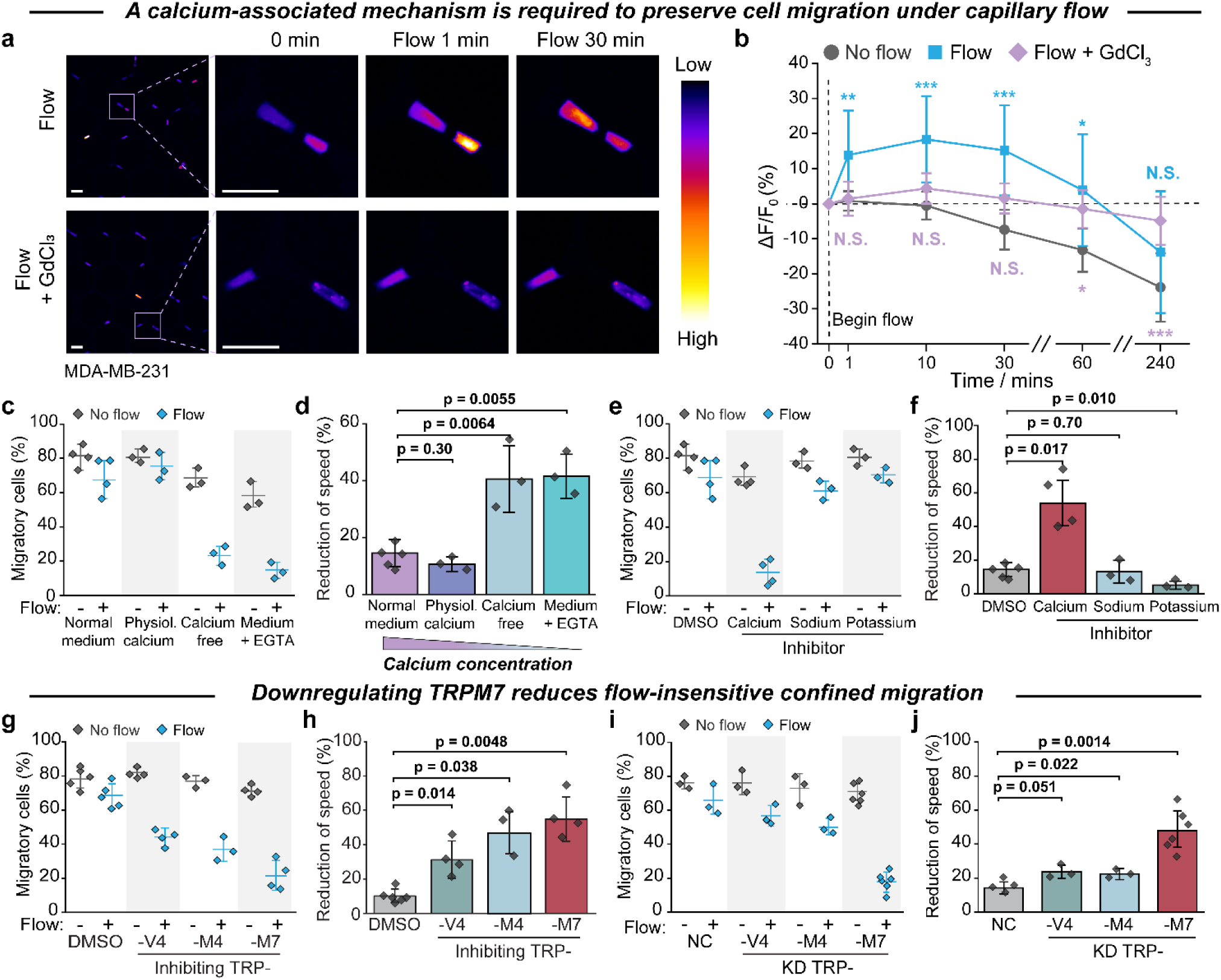
Flow-insensitive confined migration depends on the stretch-activated calcium channel TRPM7. **a and b**, Intracellular calcium dynamic in MDA-MB-231 cells after post-injection into microfluidics. Scale bars, 50 μm. **c and d**, Migration of MDA-MB-231 cells under varying external calcium environments. **e and f**, Migration of MDA-MB-231 cells treated with drugs affecting different ion channels. **g and h**, Effects of pharmacological inhibition of TRP family channels on MDA-MB-231 cell migration. **i and j**, Migration analysis of scramble negative control (NC) cells and MDA-MB-231 cells with knockdown of TRPV4, TRPM4, or TRPM7. Data are represented as the mean ± standard deviation (n ≥ 3 independent chip experiments). Statistical analysis was performed using Kruskal-Wallis tests followed by Dunn’s test (**b, d, f, h**, and **j**). P values for comparisons with the results for intracellular calcium dynamic between no flow and other two conditions are indicated (\**P* < 0.05; \*\**P* < 0.01; \*\*\**P* < 0.001). N. S., not significant.

Given the well-established roles of stretch-activated calcium channels in sensing physical cues^31^ and our finding in regulating cell migration under capillary bed-mimicking physical environment, we initially investigated Piezo1, a crucial mechanosensitive ion channel that plays a role in sensing blood shear flow^32^. Treatment of MDA-MB-231 cells with the selective peptide inhibitor GsMTx4 did not alter confined migration dynamics under either no flow and flow condition (***Extended Data Fig. 7a-b and Supplementary Video 4***). To validate this result in a more rigorous system, we conducted a Piezo1-specific knockdown in MDA-MB-231 cells and observed that their ability to perform flow-insensitive migration remained unchanged (***Extended Data Fig. 7c-e***). Accordingly, Piezo1 does not involve into this physical stresses-induced cell migration on metastatic cancer cells.

The inhibitor 2-APB, which mainly perturbs ER-associated calcium dynamics and successfully restrains cancer cell migration (***Extended Data Fig. 6d-e***), has also been found to be a universal inhibitor of mechanosensitive transient receptor potential (TRP) channels^33^. To test the potential involvement of TRP channels in this migratory behavior, we performed another pharmacological inhibition screen and found that interfering with TRPC1, TRPC4, TRPC6, and TRPV1 ion channels did not significantly influence cell migration (***Extended Data Fig. 7e-f***). However, inhibition of TRPV4, TRPM4, and TRPM7 all resulted in significant reductions in both migratory cells and migration speeds (***Fig. 2g-h***). To verify the specificity of the inhibitor screen, we performed siRNA knockdown (KD) of these three ion channels. These knockdowns did not affect cell morphology, cell migration speed on 2D culture flask, and expression of other TRP family channels (***Extended Data Fig. 7g-i***). However, TRPM7 knockdown in MDA-MB-231 cells nearly abolished motility under confinement and flow (***Fig. 2i-j* *and Supplementary Video 5***), an effect as strong as complete calcium removal from the medium (***Fig. 2c***). Additionally, increasing the extracellular concentration (∼20 mmol/L) by adding CaCl_2_ to culture medium did not restore the migratory capacity of TRPM7-knockdown MDA-MB-231 cells (***Extended Data Fig. 7j-k***). As an orthogonal approach, we activated TRPM7 channels in non-tumorigenic MCF10A cells by naltriben (NTB)^34^, but found no enhancement in cell migration under flow (***Extended Data Fig. 7l-m***). We then compared endogenous TRPM7 expression levels across a panel of breast cancer cell lines via western blot, RT-qPCR, and immunofluorescence staining (***Extended Data Fig. 8a-d***). Although previous findings that TRPM7 expression levels are elevated in several aggressive metastatic cancer samples^35^, but we found no association between TRPM7 expression and metastatic potential. Taken together, we demonstrate that TRPM7 plays a major role in regulating flow-insensitive confined migration, but as TRPM7 levels are consistent across a diverse range of cell lines, there must be an unknown TRPM7-associated factor playing a downstream role in driving this migratory behavior.

### The TRPM7-Annexin A1 signaling pathway prevents actin disassembly and facilitates migration under flow and confinement

The cytoskeleton is the main driver of cell migration dynamics, especially in response to physical stimuli^36,37^. As these dynamics have been shown to be sensitive to intracellular calcium ion concentration, we investigated the effects of cytoskeletal reorganization on flow-insensitive confined migration (***Fig. 3a-b***). While disrupting either microtubules or myosin activity had no significant effect, perturbing actin dynamics led to notable changes in migration behavior, particularly under flow condition. This was true via the inhibition of Rho-associated protein kinase (ROCK), the acceleration of actin filament depolymerization via latrunculin A, and the prevention of actin filament assembly via cytochalasin D which mediates actin cytoskeleton dynamics during cell migration and traction force generation. Actin-dependent migration occurs via lamellipodia, which are supported by the formation of branched actin networks nucleated by the actin-related protein (Arp)2/3. The use of the inhibitor CK666, which directly binds and blocks the Arp2/3 complex^38^, significantly reduced the migratory capacity of MDA-MB-231 cells under flow (***Fig. 3c-d***). On the other hand, CK869, a weaker and less selective inhibitor of the Arp2/3 complex^38^, had little effect on migration even at cytotoxic levels. Taken together, these all indicate a significant contribution of actin on flow-insensitive confined migration.

**Figure 3.**
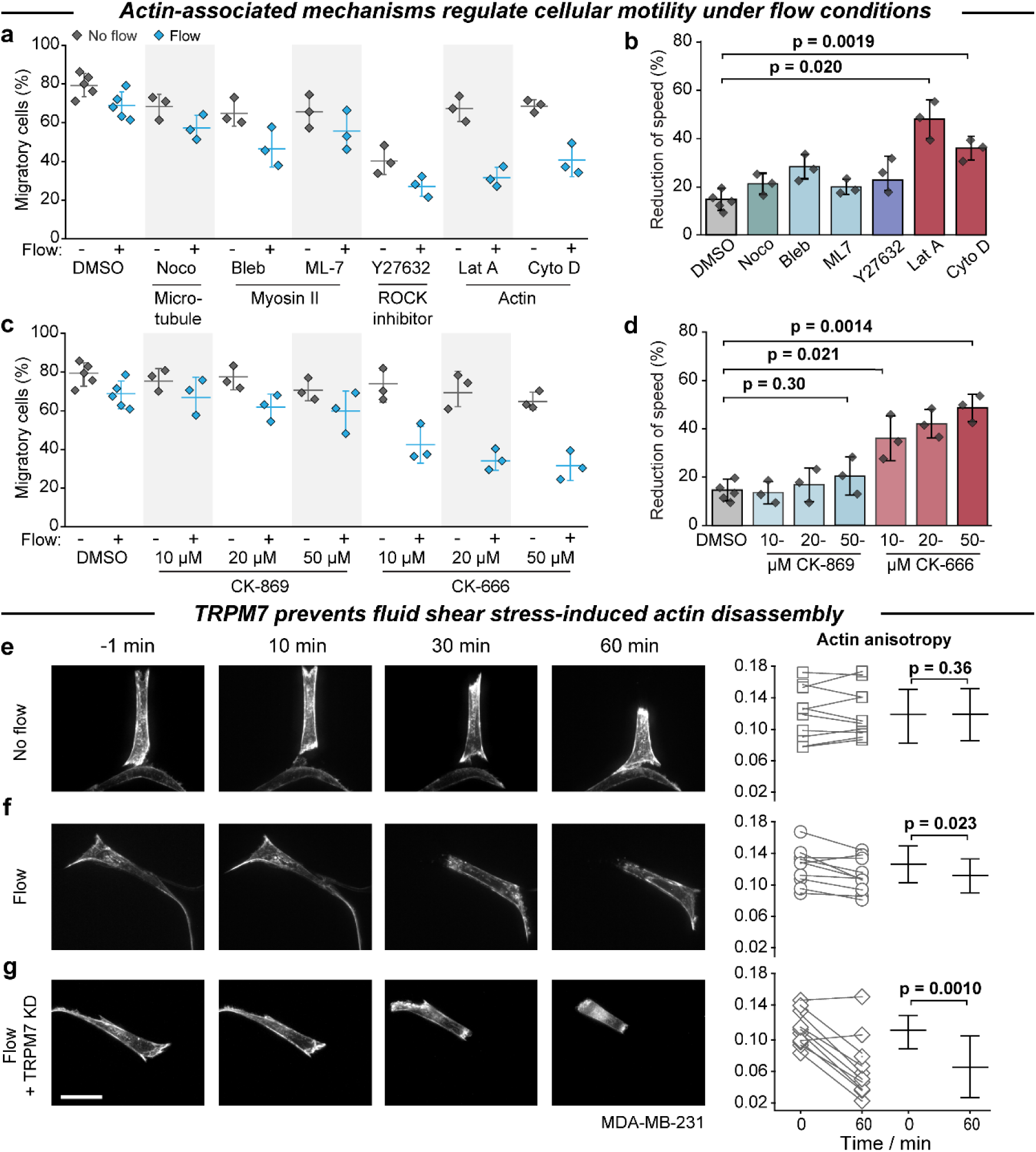
TRPM7-mediated mechanism prevents flow shear-induced cell contraction and actin disassembly. **a and b**, Effects of pharmacological drugs targeting cytoskeletal structures on MDA-MB-231 cell migration under capillary stress. **c and d**, Impact of inhibitors targeting the Arp2/3 complex on actin dynamics and their effects on MDA-MB-231 cell migration. **e, f and g**, Dynamic confocal images of Lifeact-RFP-expressing MDA-MB-231 cells and their actin anisotropy within a microfluidic chip: MDA-MB-231 cells in environments without flow (**e**), with flow (**f**), as well as TRPM7 knockdown MDA-MB-231 cells under flow conditions (**g**). Scale bars, 20 μm. Data are represented as the mean ± standard deviation (n ≥ 3 independent chip experiments). Statistical analysis was performed using Kruskal-Wallis tests followed by Dunn’s test (**b** and **d**) and one-way ANOVA followed by Tukey’s test (analysis of actin anisotropy in **e, f**, and **g**).

To better understand the connection between actin dynamic and flow-insensitive confined migration, we transfected MDA-MB-231 and MCF10A cells with Lifeact-RFP to visualize actin network dynamic. In non-metastatic MCF10A cells, initiation of flow resulted in a rapid disassembly of the actin filament network (***Extended Data Fig. 8e-f***), suggesting that MCF10A cells lose the ability to migrate due to a loss of actin filaments which is required for protrusion and contractile force formation^39^. In contrast, metastatic MDA-MB-231 cells retained their anisotropic actin filament network organization after the onset of flow (***Fig. 3e-f***). However, knocking down TRPM7 in these Lifeact-RFP cells resulted in a loss of actin filament network integrity (***Fig. 3g***), further supporting the contention that TRPM7 drives flow-insensitive confined migration via actin regulation. However, as a membrane ion channel protein, TRMP7 is not in direct contact with the actin network^34^, and thus an intermediate agent must mediate and enhance this relationship.

TRPM7 possesses a C-terminal α-kinase domain which contributes to the phosphorylation of its substrate, the signaling protein Annexin A1^40^. Annexin A1 is a Ca^2+^-dependent membrane binding protein which has been implicated in cell membrane dynamics and cytoskeleton reorganization^41^. It has been established that Annexin A1 bundles actin filaments and colocalizes with F-actin in membrane ruffles^41^. Additionally, overexpression of Annexin A1 has been shown to enhance breast cancer metastasis^42^. Therefore, our western blot analysis revealed that Annexin A1 levels are particularly elevated in breast cancer cell lines with high metastatic potential compared to non-metastatic tumor cells **(*Fig. 4a*, *Extended Data Fig. 9a-d***), despite similar TRPM7 expression levels for these breast cancer cell lines **(*Extended Data Fig. 8a-d*)**. Based on these evidences, we hypothesized that high expression level of Annexin A1 would facilitate breast cancer cell migration under flow and confinement. To test this, we first compared cell migratory capacity across multiple cell lines as a function of Annexin A1 levels, finding that breast cancer cell lines with low Annexin A1 expression can hardly migrate under flow and confinement **(*Extended Data Fig. 2b-d, and Supplementary Video 6*)**. We then generated Annexin A1 and Annexin A2 knockout 4T1 cell lines (4T1^ANXA1-/-^ and 4T1^ANXA2-/-^). Annexin A2 knockout cells exhibited major changes in cell morphology (***Extended Data Fig. 9e***), but not specifically mediating their migration under flow and confinement (***Fig. 4b-d*, *Extended Data Fig. 9f and Supplementary Video 6***), even though this protein has been shown to be a flow sensor in endothelial cells^43^. On the other hand, we found that while Annexin A1 knockout has no effect on cell morphology, it did significantly reduce the capacity of cell migration under flow environment. Furthermore, actin filaments, particularly its actin cortex structure, were highly colocalized with Annexin A1 in 4T1 cells exposed to flow (***Fig. 4e* *and Extended Data Fig. 9g***), but this colocation cannot be observed in no flow condition (***Extended Data Fig. 9h***). In contrast, flow caused a significant disassembly of actin filaments in both 67NR (low Annexin A1-level) cells and 4T1^ANXA1-/-^ cells (***Extended Data Fig. 9g-h***), highlighting the critical role of Annexin A1 in maintaining actin structure under flow. Together, these results show that Annexin A1 plays a key role in enabling flow-insensitive confined migration. As a unique mechanism for highly metastatic cancer cells, those metastatic breast cancer cells utilize a TRPM7-Annexin A1 to prevent actin filament disassembly, allowing for sustained cancer cell migration under capillary flow and confinement.

**Figure 4.**
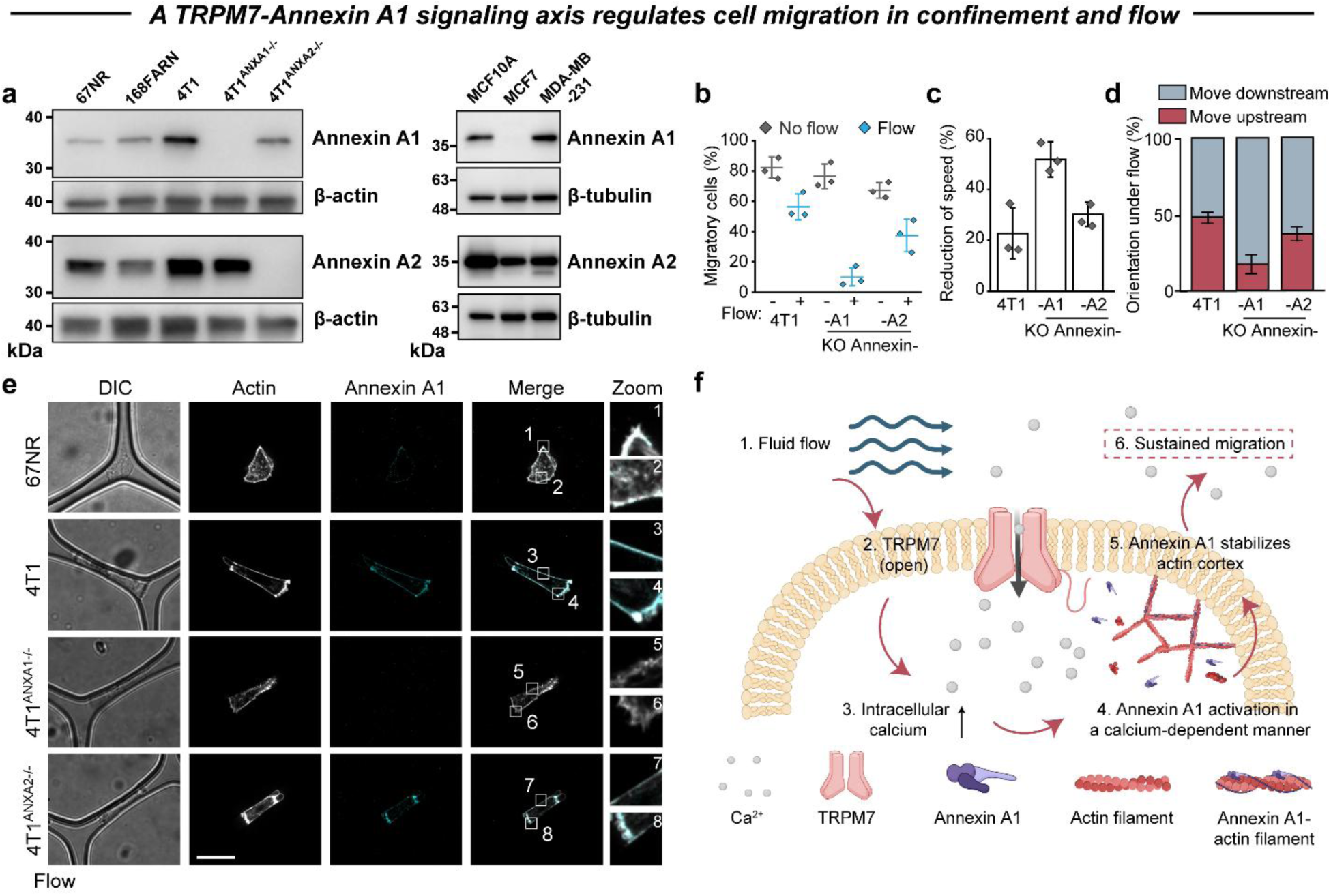
High expression level of Annexin A1 on metastatic breast cancer cells prevents cytoskeleton reorganization, ensuring cell migration under flow condition. **a**, Western blot analysis of Annexin A1 and Annexin A2 expression levels in breast cancer cells. **b-d**, The influence of knockout Annexin A1 and Annexin A2 on cancer cell migration. **e**, Immunofluorescence analysis showing the co-localization of Annexin A1 and F-actin under flow conditions. Scale bars, 20 μm. **f**, Schematic of the proposed mechanotransduction pathway in metastatic breast cancer cells responding to flow and confinement. Data are represented as the mean ± standard deviation (n ≥ 3 independent chip experiments).

To determine the *in vivo* relevance of the newly identified TRPM7-Annexin A1 mechanosensitive signaling pathway in promoting cell migration under capillary vessels, we established an intravital imaging system using two-photon synthetic aperture microscopy^18^, which offers low phototoxicity for long-term monitoring of *in vivo* metastasis (***Fig. 5a***). MDA-MB-231 cells were labeled with CellTracker Deep Red and resuspended prior to injection into the spleen of C57BL/J mice. To monitor the metastatic process of the injected circulating tumor cells, we exposed and imaged the liver, which is anatomically connected to the spleen via the portal vein system^13^ (***Extended Data Fig. 10a***). Once individual cancer cells were trapped in the microvasculature of the liver, they resisted dispersal under flow unless cellular rupture occurred (***Extended Data Fig. 10b***). A portion of the injected MDA-MB-231 cells remained attached to the liver vasculature and did not move, while others migrated along capillary vessels **(*Fig. 5b* *and Supplementary Video 7***). In contrast, either TRPM7 knockdown cells or Annexin A1 knockdown cells significantly block their cell migration in capillary bed (***Fig. 5c-e*)**. By comparing their morphological reorganization and protrusion dynamics, we observed that reduced levels of either TRPM7 or Annexin A1 significantly decreased the ability of cell to reorganize cell morphology within the capillary bed *in vivo* (***Fig. 5b-d*)**, a process that is heavily dependent on actin cytoskeleton dynamics^39^. These *in vivo* results confirm the powerful role of the mechanosensitive TRPM7-Annexin A1-actin signaling pathway in mediating the dissemination of circulating cancer cells during metastasis (***Fig. 4f***).

**Figure 5.**
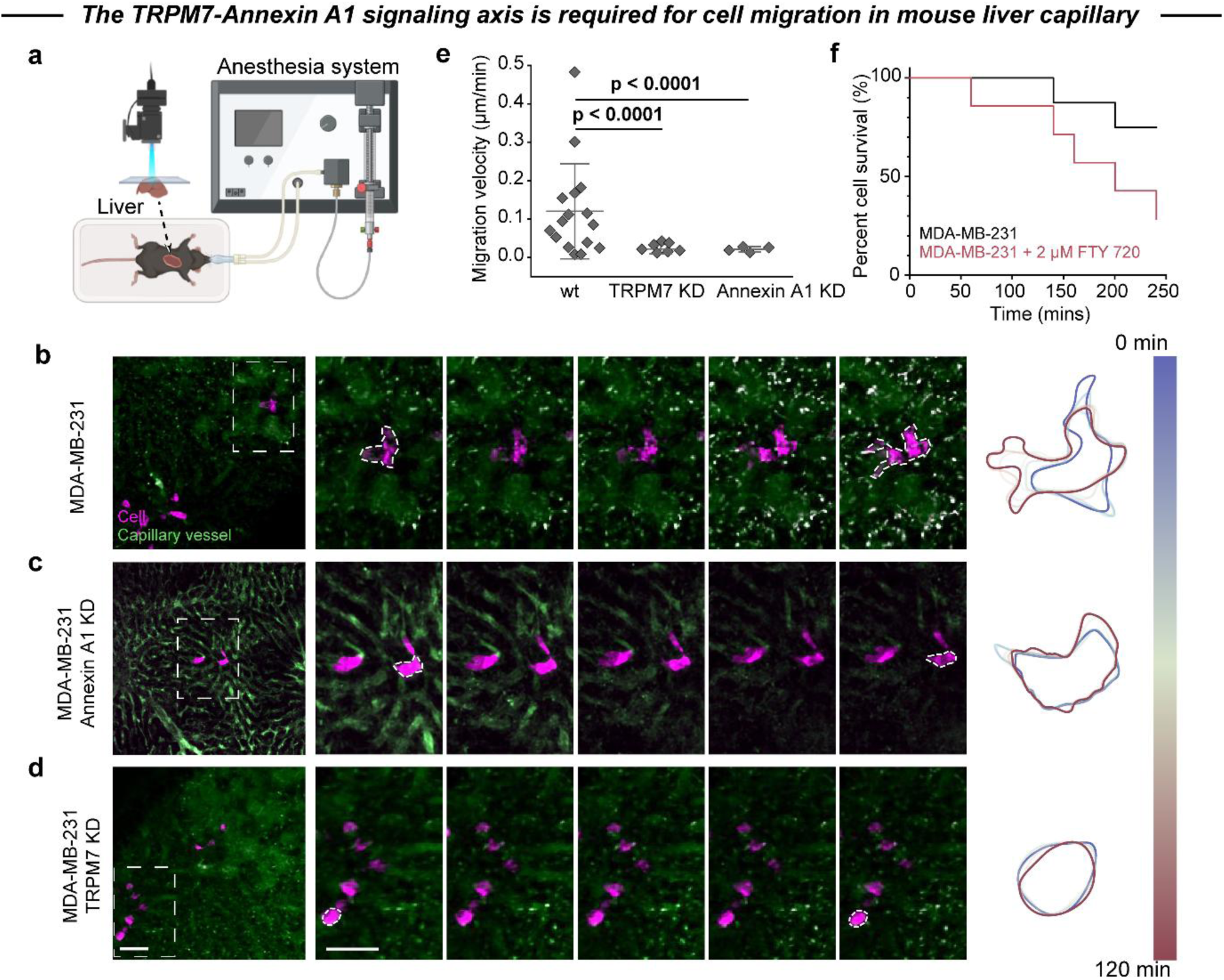
The TRPM7-Annexin A1 signaling axis in metastatic breast cancer cells facilitates migration within liver capillary vessels. **a.** Illustrations of the imaging setup**. b-d**, Two-photon intravital imaging to track breast cancer cell migration within mouse liver capillaries. **e**, Analysis of cell migration velocity within capillary vessels. **f**, Survival curve of MDA-MB-231 in mouse liver with/without FTY720 treatment. Scale bars, 50 μm. Data are represented as the mean ± standard deviation. Statistical analysis was performed using one-way ANOVA followed by Tukey’s test (**e**).

### The immunomodulatory drug FTY720 efficiently and selectively eliminates breast cancer cells under flow and confinement

Building on our *in vitro* and *in vivo* results showing the key role of TRPM7 in mediating metastatic dissemination of cancer cells through capillary bed, we sought to identify a pharmaceutical approach to target this TRPM7-based mechanosensing. We found FTY720 (Fingolimod), an FDA-approved immunomodulatory drug currently used to treat multiple sclerosis, based on separate *in vitro* and *in vivo* studies that have shown it to be a TRPM7 inhibitor^44,45^. This FTY720 treatment highly reduced the cell viability of metastatic MDA-MB-231 cells only under confinement and flow as evidenced by caspase 3 staining (***Fig. 6a-d*, *Extended Data Fig. 11a-b, and Supplementary Video 8***). Interestingly, and importantly in the context of the requirements for anti-cancer drugs to be cancer cell-specific, we found that FTY720 treatment of non-tumorigenic MCF10A cells did not result in significant levels of cell death (***Fig. 6e-f*, *and Extended Data Fig. 11c-d***). The positive control chemotherapy drug doxorubicin was found to kill all types of cells in all physical conditions in a nonspecific fashion (***Extended Data Fig. 11e-h***). However, some calcium-associated inhibitors, such as GdCl_3_ and 2-APB, which effectively restricted cell migration, did not actually induce cell rupture in cancer cells (***Extended Data Fig. 11i-j***).

**Figure 6.**
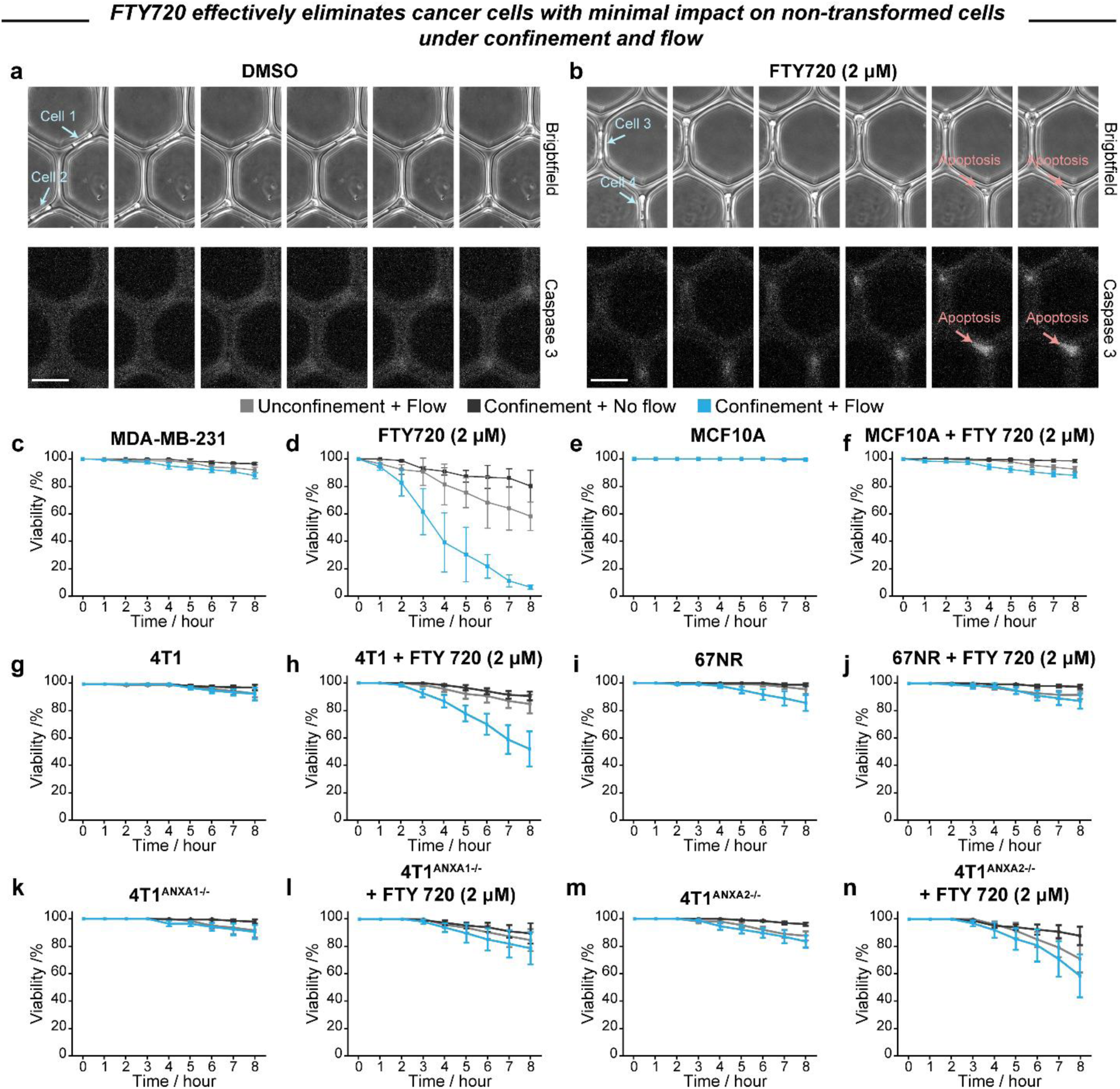
Potential mechano-pharmacology of FTY720 and its lethal effect on breast cancer cells under flow and confinement. **a and b**, Caspase-based method for identifying cell death and rupture in a capillary microfluidic chip. The interval time between each time-lapse image is 30 minutes. Scale bars, 50 μm. **c-f**, FTY720 induces significant cell rupture in highly metastatic MDA-MB-231 cells but has minimal impact on the survival of non-tumorigenic MCF10A cells. **g-j**, FTY720 similarly increases cell death in metastatic mouse mammary carcinoma 4T1 cells but does less affect non-metastatic mouse mammary carcinoma 67NR cells. **k-n**, Knockout of Annexin A1, but not Annexin A2, reduces the sensitivity of FTY720-associated rupture in 4T1 cells. Data are represented as the mean ± standard deviation (n ≥ 3 independent chip experiments).

To better understand this metastatic cell type-specific, FTY720-induced cell rupture, we assessed the viability of TRPM7-KD MDA-MB-231 cells following FTY720 treatment. To gain deeper insights into the metastatic cell type-specific cell death induced by FTY720, we evaluated the viability of TRPM7-knockdown MDA-MB-231 cells after FTY720 treatment. Although these cells are much less motile in environments presenting confinement and flow, they are still killed efficiently when exposed to FTY720 (***Extended Data Fig. 11k-l***). Although these cells exhibit significantly reduced motility under conditions of confinement and flow, they remain highly susceptible to FTY720-induced cell death when exposed to confinement and flow. This observation indicates that FTY720’s mechanism of action likely operates through downstream effectors of TRPM7. To test this, we examined the impact of FTY720 on non-metastatic 67NR cells and metastatic 4T1 cells. Reflecting our findings in MDA-MB-231 cells, we found that metastatic 4T1 cells were killed at a much higher level in response to FTY720, than non-metastatic 67NR cells. (***Fig. 6g-j* *and Supplementary Video 9***) Consistent with our observations in MDA-MB-231 cells, we discovered that metastatic 4T1 cells exhibited significantly higher levels of cell death in response to FTY720, confinement, and flow compared to non-metastatic 67NR cells. As we earlier found Annexin A1 to be overexpressed in 4T1 cells compared to 67NR cells (***Extended Data Fig. 9a***), we looked to test the effects of FTY720 on cells with controlled Annexin levels. In 4T1^ANXA1-/-^ cells exposed to FTY720, confinement, and flow, minimal cell death was observed. In contrast, 4T1^ANXA2-/-^ cells exhibited significant levels of cell rupture under the same conditions (***Fig. 6k-n***).

Finally, to determine if this effect was also present *in vivo*, we injected both wild type and FTY720-treated MDA-MB-231 cells into mice as previously described. Over a 4-hour intravital microscopy observation period, more than 2.5 times as many control cells survived compared to FTY720-treated cells, which exhibited frequent rupture (***Fig. 5f***). This demonstrates that the cytotoxic effects from FTY720 on metastatic cancer cells are also effective under *in vivo* microenvironments. Taken together, these results show that FTY720 possesses cytotoxic effects in cells exhibiting Annexin A1 overexpression that are exposed to capillary physical cues. Although FSS-induced cell rupture has been reported in previous *in vitro* and *in vivo* studies^9,12,13,46,47^, this is the first observation of a drug with the ability to efficiently and selectively kill cancer cells in a mechanosensitive FSS-dependent fashion as they traverse the blood vessels required for metastasis.

## Discussion

Collectively, our findings elucidate a TRPM7-Annexin A1-mediated mechanism that enables metastatic breast cancer cells to resist FSS-induced actin filament disassembly, thereby facilitating their confined migration within the bloodstream (***Fig. 4f***). In the context of cancer metastasis, this tailored mechanism may help cancer cells overcome the physical entrapment during circulation and promote subsequent metastatic processes, such as active extravasation, ultimately allowing them to become part of the 0.1% of tumor cells that successfully metastasize. This finding also provides an explanation for why the knockdown of TRPM7 in highly metastatic MDA-MB-231 cells led to a loss of metastatic capacity in mouse models, while not impacting cellular proliferation, as observed in previous research^48^.

Notably, successful metastasis requires tumor cells to tailor their cellular behaviors. These tailored features in response to mechanical cues may also represent potential vulnerabilities that can be exploited for therapeutic intervention. In this study, we introduce a novel mechano-pharmacology targeted to metastatic cancer cells in circulation by FTY720. While FTY720 is primarily used for its immunomodulatory effects in treating multiple sclerosis, previous studies have demonstrated its anti-cancer potential^49,50^, particularly against triple-negative breast cancer. Our findings further reveal that this approach may induce extensive cancer cell rupture under the physical stress of blood flow. Only cancer cells, but not non-transformed cells, become sensitive to FTY720-induced rupture under microcirculation. This phenomenon is a mechanics-driven strategy but provide high efficiency and specificity for eliminate the cancer cells. Beyond this work, a deeper understanding of mechanobiology and cancer metastasis may help identify more precise and safer mechano-pharmacology biomarkers, ultimately reducing or even eliminating systemic adverse effects in cancer therapy.

## Methods

### Cells and cell culture

Human breast epithelial cancer cell lines, MCF7, and MDA-MB-231, and immortalized human breast epithelial cell line MCF10A were all purchased from ATCC. Murine breast cancer cell lines 67NR, 168FARN, and 4T1 were gifts from Hsih Kim, Lina Lim’s group. Briefly, the 67NR, 168FARN and 4T1 are isogenic cell lines isolated from a spontaneously arising tumor and separated based on key phenotypic characteristics. 67NR cells are non-invasive and non-metastatic, 168FARN cells are invasive but non-metastatic while 4T1 cells are both invasive and metastatic. Thus, these three cell lines represent different levels of metastatic potential but identical genetic origins^51^. All breast epithelial cancer cell lines were cultured in high glucose DMEM (Lonza) supplemented with 10% FBS (Gibco) and 5 μg/ml gentamycin (Gibco). MCF10A cell line was cultured in MEBM (Lonza) supplemented with MEGM kit (Lonza). HCC38 was a gift from G.V. Shivashankar’s group and BT549 was a gift from George Yip’s group, which were purchased from ATCC. These two cell lines were cultured in RPMI 1640 (Lonza) supplemented with 10% FBS (Gibco) and 5 μg/ml gentamycin (Gibco). To reduce the potential side effects caused for long-phase growth, all cell lines used in this study would maintain a lower subculture number (less than 10 passage times). All cell lines used in this study were maintained in tissue culture flash (SPL) at 37°C with 5% CO_2_ and passed weekly. Medium refreshing was executed every two days to keep the nutrition for cell growth. Routine mycoplasma contamination has been tested at least twice each year by using either MycoAlert PLUS mycoplasma contamination kit (Lonza) or MycoSEQ PCR kit (Thermofisher). Before experiments, all cells grew to around 90%∼95% confluency and changed cell culture medium 12 hours ahead. During the experiment, adherent cells were washed by DPBS x 2 times and detached by incubating with TrypLE (Gibco) for 10 minutes (12 mins for MCF10A cell) in a 37 °C incubator. After spinning down (1000 RCF for 3 mins), the cell pellets were collected and resuspended in fresh complete medium at a concentration of 0.8∼1 × 10^6^ cells/ml for the subsequent microfluidic experiment.

### Fabrication of microfluidic devices

Standard one-layer soft lithography was performed to create the polydimethylsiloxane-based microfluidic device consisting of an array of microchannels with different morphology features (height: 20 μm). In brief, a 20 μm thick layer of SU8 photoresist was spincoated onto a silicon wafer and crosslinked with UV light (μFabrication service was provided by the microfabrication core facility in Mechanobiology Institute). The master wafer surfaces were treated with oxygen plasma at 15W, 8.8 sccm for 30s (Tergeo, PIE Scientific), and silanized (1H, 1H, 2H, 2H-perfluorooctyle trichlorosilane, Cat# 448931, Sigma-Aldrich) for 2 hours under vacuum. The mixed-well polydimethylsiloxane (PDMS, Sylgard 184 Elastomer Kit, Dow Corning) at a ratio of 10:1 covered the silanized wafer and degassed for half hour to remove the bubbles. Then, the devices were placed in a thermal oven (Memmert) at 70 °C for at least 6 hours bake (but no more than 8 hour). After taking out and cooling down to room temperature, a scalpel was employed to peel off the mold and cut it into square chips. Next, the PDMS chips were punched with 1.5 mm holes for creating inlets and outlets and bonded to a glass culture dish (27mm IWAKI glass dish or Cellvis 27mm glass dish) after oxygen plasma (15W, 8.8 sccm for 20s) activating. The final devices were placed in a 70 °C oven for one hour baking to facilitate irreversible binding. The chips were subjected to UV for 30 minutes inside a biosafety cabinet. A serum-free cell culture medium solution with collagen IV (20 μg/ml, Sigma-Aldrich, C0543) was injected into microfluidic devices and incubated overnight at 4 °C. Before the experiment (usually one hour ahead), the solution inside microfluidic devices would be replaced to complete medium.

### Microfluidic assay for testing cell migration

An illustration for brief description has been shown in Extended Data Fig. 1e. Microfluidic chips were coated with ECM proteins (50 μg/ml COL type IV, except for the experiment to compare the effect of different ECM proteins) one day ahead. Following the cell culture process, the suspended cells were loaded into a 1 ml syringe (NIPRO) with a UNP-23 precision tip (Unicontrols) connected to a Tygon tubing (0.02 inch inner and 0.06 mm inch outer, Cole-Parmer). The medium with suspended cells was injected through the microchannels at a volume flow rate of 0.7 μL/min for 10 minutes via a syringe pump (Fusion 200, CHEMTX Inc.). At the end of the injection process, the tube was snipped at an appropriate length by using a scissor. This residual tube would function as an outlet for effluent collection in the following experiment (**Extended Data Fig. 2**). All the processes must avoid any generations of air bubbles.

Subsequently, the microfluidic chips with captured cells were immediately transferred to culture incubator or IMQ Biostation (Nikon) with 37°C and 5% CO_2_ humidified chamber for dynamically tracking the cell behaviors. The imaging interval was 5 mins for 24 hours and images were taken using the 10x and 20x ph objective. A syringe-based system was connected to the inlet hole and the constant flow was created by the syringe pump. The pump volumetric flow rate was set at 0.15 μL/min by following the results in the flow optimization section. Live-cell videos were exported to ImageJ (National Institutes of Health) and the ‘Manual Tracking’ function was used to analyze cell motility, including migratory cells, migration speed, migration path, and migration orientation. ‘Manual Tracking’ was also applied to measure cell viability.

### Generation of liquid flow speed via microbead assay and COMSOL simulation in microfluidic device

For creating a capillary-mimicking mechanical microenvironment, the liquid flow generated by syringe pump must reach a physiologically relevant flow rate. Microbeads were injected into chip device for measuring the flow rate. Briefly, the 2 μm microbeads (Invitrogen, F13838) were diluted into DMEM as 0.5% *wt* concentration and injected into microfluidic device. The flowing process was recorded by a high-speed camera (FASTCAM Mini AX50, MindVision, Japan) under room temperature. The average flow rate of microbeads was analyzed by using the manual tracking function in ImageJ (National Institutes of Health).

The flow rate of fluids in microfluidic device was simulated by the COMSOL 5.4. In brief, a laminar flow-based method was applied to calculate the flow rate in confined channel area, where the model simplifies as steady state to mimic the long-term flow expose. The volume flow rate applied to the simulation was based on the parameter (also volume flow rate) provided by the syringe pump system. Because of the present of tributaries, the results only provide the range of max and min flow rate in confined channel but not the average flow rate.

### Theoretical calculation of shear flow in microfluidic device

During metastasis in microcirculation, the arresting CTC usually must stay on a constricted capillary vessel with blood liquid pressure and FSS. The previous works found that exposing to high FSS can destroy the CTC in a microfluidic system^12,52,53^. For further study, the measurement of FSS should be covered.

In brief, we applied Navier-Stokes equation under Newtonian fluids and laminar flow to model the FSS in microfluidic device. While the exact solution for flow in rectangular ducts describe as a wall shear stress (τ_w_). The relationship are derived for pressure drop and the product of Reynolds number and Fanning friction factor, fRe, which is calculated by using following formula^54^.

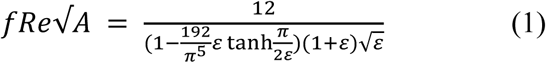

ε presented in terms of the aspect ratio of cross-section, 0≤ε≡c/b≤1, where b and c are the major and minor semi-axes of the cross-section, b≥c. In our chip design, the value of ε reach 0.25∼0.5, where was calculated as 16.48∼22.77.

The Fanning friction factor is defined as:

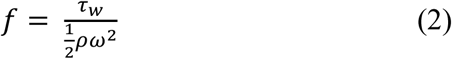

where ρ is the density of the liquid and ω is the flow velocity.

And the square root of area-associated Reynolds number is defined as:

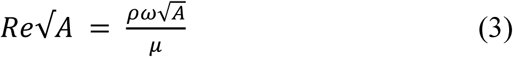

where μ is the dynamic viscosity of the fluid, considering as 0.01 N·s/m^2^ in culture medium. Combining Eqs. (2) and (3) becomes:

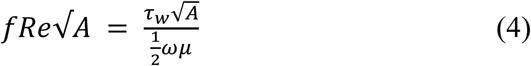

And rewrite as:

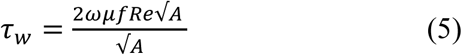

Based on the number we acquired above, we identified the FSS in our microfluidics device is 5.9∼11.8 N/m^2^, where is similar to the FSS (typically around 5∼40 N/m^2^).

### ECM coating

To compare the effects of different ECM proteins, microfluidic chips were coated with 50 μg/ml collagen type I (COL I, Corning, 354236), 50 μg/ml collagen type IV (COL IV, Sigma, C0543), 20 μg/ml fibronectin (Sigma, F0895), or 1% F-127 (vol/vol, Invitrogen, P6866) in DMEM (/RPMI 1640/MEGM) without FBS overnight in 37 °C incubator. Before experimentation, the coating solution was replaced with fresh growth medium (DMEM/RPMI 1640/MEGM).

### Immunofluorescence

In this study, we imaged the cells after 8-hour post-injection. For imaging the cells in the microfluidic chip, 10 μL of 4% formaldehyde solution (Sigma-Aldrich) was injected into the channel with 0.3 μL/min. The fixing process was quenched with DPBS containing 100 mM glycine. Then cells were permeabilized by using DPBS with 0.5% Triton X-100 (Sigma-Aldrich) and blocked against nonspecific binding with DPBS containing 0.1% Triton X-100 and 2.5% bovine serum albumin (BSA, Sigma-Aldrich). Next, the primary antibody (Paxillin, ab32084, Abcam, 1:250; TRPM7, ab262909, Abcam, 1:500; Annexin A1, ab65844, Abcam, 1:500) was diluted in antibody dilution buffer (DPBS containing 0.1% Tween-20 and 1% BSA) and injected into a microfluidics chip for overnight incubation in a 4 °C fridge. Later, the cells were incubated with secondary antibodies (1:1000, Alexa Fluor 488 and 594 goat anti-mouse and/or anti-rabbit IgG, Abcam) diluted with antibody dilution buffer at room temperature for 1 hour. For nucleus and actin labeling, cells were stained with Hoechst 33342 (1:1000, 10 mg/ml, H3570, Invitrogen) and Alexa Fluor 647-phalloidin (1:2000, A22287, Invitrogen). After the staining process, cells were washed with 10 μL 0.1% DPBS-T. All the immunofluorescence images presented in this study were taken by a W1 confocal microscope (Nikon) with the 100x oil objective.

For imaging the suspended cells in plate, glass bottom dish should be incubated by DPBS with 1% Pluronic F-127 (Invitrogen) for 1 hour. Later, gently washing by DPBS. Cell pellets caught from cell culture process incubates with 4% formaldehyde solution (Sigma-Aldrich) for fixing the cell within 20 mins. After fixing, cell pellets should be spun down using a benchtop microcentrifuge at 6000 rpm for 30 s and re-suspended into DPBS containing 100 mM glycine for quenching the fixing process. Next, cell pellets were transferred into the glass bottom dish with anti-adhesive surface and spun down within 1 hour. Then cells were permeabilized by using DPBS with 0.5% Triton X-100 (Sigma-Aldrich) and blocked against nonspecific binding with DPBS containing 0.1% Triton X-100 and 2.5% bovine serum albumin (BSA, Sigma-Aldrich). For nucleus and actin labeling, cells were stained with Hoechst 33342 (1:1000, 10 mg/ml, H3570, Invitrogen) and Alexa Fluor 647-phalloidin (1:2000, A22287, Invitrogen). After the staining process, cells were washed with 0.1% DPBS-T for three times. All the pipetting process should be gently to reduce the losing of suspended cells. All the immunofluorescence images presented in this study were taken by a W1 confocal microscope (Nikon) with the 100x oil objective.

### Cell volume and nuclear volume measurement

Cell volume was measured by Alexa Fluor 647-phalloidin and nuclear volume was measured by Hoechst 33342. In briefly, the fixing cells were followed the IF stain protocol introduced above and stained actin cytoskeleton with phalloidin and nucleus with Hoechst 33342. Then, the stained-well cells were placed on a W1 confocal microscope and imaged with a 100x oil objective at Z-position (0.5 μm/layer, ±12 μm). The 3D confocal microscopy images were analyzed by using Imaris (version 9.6) image analysis software (Bitplane).

### Artificial sort of nuclear subpopulation via flow cytometry

To sort the cells with small nucleus, we employ the flow cytometry for doing a cell size-based sort. In briefly, the suspending cells at a concentration of around 5 million cells/ml were passed through a strainer with 40 μm hole to remove the cell cluster. A Nikon flow cytometry (Nikon SH800) was employed to sort the cells with different nucleus range. After finishing sorting process, those sorted cells were immediately injected into capillary chip and observed their migratory features under biostation live cell imaging system.

### Pharmacological treatments

In this study, pharmacological inhibitors were used, including: 1 mM EGTA (Sigma-Aldrich) chelated calcium in culture medium, 10 μM Gadolinium (III) chloride (Gd^3+^) (Tocris Bioscience) inhibited calcium channels, 2 μM phenamil methanesulfonate (Sigma-Aldrich) inhibited sodium channels, 5 μM UCL2077 (Sigma-Aldrich) inhibited potassium channels, 10 μM JIB04 (Sigma-Aldrich), a histone demethylase inhibitor increases the levels of nucleus condensation, and 10 μM RGFP966 (Sigma-Aldrich), A histone deacetylases 3 inhibitor results in less condensation of cell nucleus, 10 μM BAPTA-AM (Sigma-Aldrich) chelated intracellular, 100 μM 2-APB interfered calcium release from ER, 20 μM sodium deoxycholate (DOCL), 5 μM GsMTx4 (Tocris Bioscience) affects Piezo 1 mechanosensitive ion channel on the plasma membrane, 0.2 μM Pico145 interfered TRPC 1 channel, 5 μM ML204 (Sigma-Aldrich) interfered TRPC 4, 1 μM SAR7334 (Sigma-Aldrich) interfered TRPC 6, 10 μM SB366791 (Sigma-Aldrich) interfered TRPV 1, 20 μM RN1747 (Sigma-Aldrich) interfered TRPV 4, 20 μM 9-Phenanthrol (Sigma-Aldrich) interfered TRPM 4, 2 μM NS8593 (Tocris Bioscience) interfered TRPM 7, 5 μM Naltriben (Sigma-Aldrich) activated TRPM 7, 5 μM Nocodazo (Noco, Tocris Bioscience) increase the depolymerization of microtuble, 10 μM myosin II ATPase inhibitor Blebbistain (Bleb, Tocris Bioscience), 20 μM myosin light chain kinase inhibitor ML-7 (Tocris Bioscience), 10 μM Rho-associated protein kinase (ROCKs) inhibitor Y27632 (Tocris Bioscience), 0.5 μM Latrunculin A (Lat A, Santa Cruz Biotechnology Inc., SCB Inc.) disrupts actin filament dynamics and prevent G-actin polymerization into filaments, 1 μM Cytochalasin D (Cyto D, Sigma-Aldrich) disrupts actin polymerization by binding to the barbed ends of actin filaments, 10, 20, and 50 μM CK-869 (Sigma-Aldrich) and CK-666 (Sigma-Aldrich) target the actin-related protein 2/3 and inhibit actin polymerization, but the inhibition mechanism is different. 0.2, 2, and 20 μM FTY 720 (Sigma-Aldrich) targets to sphingosine-1-phosphate receptor. A gradient of doxorubicin (DOX, Sigma-Aldrich), a traditional chemotherapy drug, disrupts the normal structure of DNA. All reagents were dissolved into DMSO (Sigma-Aldrich). To minimize the side effects caused by DMSO (control group), all pharmacological solutions were pre-prepared at 1000× the experimental concentration. In all pharmacological inhibition experiments, the medium was supplemented with 0.1% DMSO (vol/vol) or 0.1% DMSO (vol/vol) with inhibitor, forming mixing solution. In pharmacological treatment experiment, fresh complete medium (**Fabrication of microfluidic devices**) has been replaced as the mixing solution to wash microfluidic chips. Cell pellets were resuspended into the mixing solution and injected into microfluidics immediately.

### Calcium concentration in growth medium

The calcium concentration obtained from the Lonza company is around 7∼8 mmol/L in DMEM. Calcium concentration in FBS is around 2.7∼3.1 mmol/L. Physiologically related calcium concentration in healthy human blood 2.1∼2.6 mmol/L. Calcium free DMEM with 10% FBS (Calcium concentration: 0.24∼0.28 mmol/L) and normal DMEM (Calcium concentration: 6∼7 mmol/L) was mixed in a certain proportion to obtain a mixing culture medium with physiologically related calcium concentration for microfluidic experiments. To obtain the culture medium with higher calcium concentration (∼20 mmol/L), calcium chloride (Sigma) has been introduced into normal DMEM.

### Intracellular calcium imaging

To visualize the concentration of intracellular calcium, we employed Fluo-4 Am Direct Calcium Assay Kit (ThermoFisher). In brief, 2x working buffer was prepared based on the manufacturer’s protocol. And the working buffer was diluted 1x with culture mediate and incubated with suspended cells for 20 mins at 37°C and 5% CO_2_. Subsequently, Fluo-4 Am buffer solution was removed by spin down (200 rpm, 1min) and the cell pellets were re-suspended into fresh culture medium. Following the microfluidic protocol, cells were injected into microfluidic chip and immediately placed on a W1 confocal microscope (Nikon) -- living cell imaging system, which was equipped with a semi-enclosed incubating system maintained at 37 °C and 5% CO_2_. Cells were imaged with Z-position (1 μm/layer, ± 2 μm) in 0, 10, 30, 60 mins. Then the intervals set would change to every 60 mins for next 5 hours under flow (or static) conditions. Dynamic intracellular calcium was recorded by GFP signal. The fluorescent intensity of individual cells was analyzed using ImageJ (National Institutes of Health). Z project in static function is requested to minimize the error caused by the difference of cell height and the fluorescent intensity for each cell was quantified by outlining the individual cell using the polygonal tool and measuring by analyzing function.

### Quantitative PCR analysis

To verify the gene experiment level of all breast cancer cells, an RNA extraction kit (QIAGEN, RNeasy Plus Mini Kit) was employed to extract the RNA from cells when they grew to around 90∼95% of confluence in a T25 cell culture flash. Each group was done in three duplications. The extracted RNA was transferred to cDNA (Bio-Rad, iScript Adv cDNA Kit for RT-qPCR) and stored on 4 °C fridge but no longer than one week. All the primers were purchased from Integrated DNA Technologies (IDT), including: GAPDH: F-(5’-CAAGCTCATTTCCTGGTATGAC- 3’) and R-(5’-CAGTGAGGGTCTCTCTCTTCCT-3’), UBB: F-(5’-GCTTTGTTGGGTGAGCTTGT-3’) and R-(5’- CGAAGATCTGCATTTTGACCT -3’),Piezo 1: F-(5’-TTCCTGCTGTACCAGTACCT-3’) and R-(5’- AGGTACAGCCACTTGATGAG -3’), TRPC 1: F-(5’- GATGTGCTTGGGAGAAATGC-3’) and R-(5’- AATGACAGGTGCAACATCCA-3’), TRPV 4: F- (5’-CCCGTGAGAACACCAAGTTT-3’) and R-(5’- GTGTCCTCATCCGTCACCTC -3’), TRPM 4: F-(5’-TGCGCGCCGAGATGTAT-3’) and R-(5’- AAAGAAGCAGGTCGCTCCAG -3’), TRPM 7: F- (5’-GGAGATGCCCTCAAAGAACA-3’) and R-(5’- TGCTCAGGGGGTTCAATAAG-3’).

The preparation of SYBR Green based-cocktail solution (SYBR Green Master Mix, Bio-Rad) for RT-qPCR was following the protocol in our previous work^55^. The mixed well PCR cocktail was subsequently placed on Bio-Rad CFX96 (Bio-Rad) after centrifuging (200 rpm, 1 min). The following thermal setting was applied on the RT-qPCR cycler: 95°C for 10 min followed by 40 cycles of amplification (95°C for 20 s, 55°C for 30 s, and then 72°C for 20 s) and final additional incubation at 72°C for 5 mins. The relatively gene expression data were normalized to reference gene (GAPDH and UBB) with the following equation:

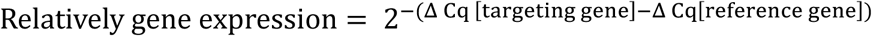

### siRNA knockdown experiment

To generate MDA-MB-231 cells with special gene knockdown, RNAi (all DsiRNA sequences purchased from IDT) for targeting gene knockdown or scramble control RNA was transfected (jetPrime, Polyplus Transfection) for 24 hours in 70% confluent cells seeded in a 6 well culture plate. Two target sequences were generated for TRPM7 and found to be equally effective at reducing TRPM7 mRNA levels (**Extended Data Fig. 7h**). Both TRPM7 target sequences induce MDA-MB-231 restrain their migratory capacity under flow environment. And TRPM7_1 sequence performed stronger inhibition for cell migration. All subsequent TRPM7 knockdown experiments were performed using the TRPM7_1 sequence. The sequences for targeting genes are as follows:

Scramble negative control sh

(5’-rCrUrUrCrCrUrCrUrCrUrUrUrCrUrCrUrCrCrCrUrUrGrUGA-3’)

(3’-rUrCrArCrArArGrGrGrArGrArGrArArArGrArGrArGrGrArArGrGrA-5’)

TRPM4

(5’-rGrArGrArArGrArUrGrUrUrGrArCrGrCrGrArArUrArGrAGA-3’)

(3’-rUrCrUrCrUrArUrUrCrGrCrGrUrCrArArCrArUrCrUrUrCrUrCrArU-5’)

TRPV4

(5’-rGrGrCrGrArGrGrUrCrArUrUrArCrGrCrUrCrUrUrCrArCTG-3’)

(3’-rCrArGrUrGrArArGrArGrCrGrUrArArUrGrArCrCrUrCrGrCrCrArG-5’)

TRPM7_1

(5’-rGrUrGrArArGrArArUrCrArArUrGrGrCrUrArArArGrCrATT-3’)

(3’-rArArUrGrCrUrUrUrArGrCrCrArUrUrGrArUrUrCrUrUrCrArCrCrA-5’)

TRPM7_2

(5’-rCrArArArCrCrUrGrArArGrUrCrArUrUrCrUrGrCrArArCTT-3’)

(3’-rArArGrUrUrGrCrArGrArArUrGrArCrUrUrCrArGrGrUrUrUrGrGrU-5’)

The knockdown efficiency was verified and optimized via RT-qPCR method. Through optimization, the knockdown cocktail was fine-tuned to achieve a silence efficiency of over 80% for the target gene.

### Generation of ANXA1 and ANXA2 knock-out cell lines

ANXA1 CRISPR/Cas9 KO (sc-421459) and homology-directed repair (sc-421459-HDR) plasmid constructs were obtained from Santa Cruz Biotechnology (SCBT), and co-transfected into 4T1 cells using Lipofectamine^TM^ 3000 reagent (Invitrogen). 48 hours post transfection, cells were selected using puromycin at 5 µg/mL for 3 days. Subsequently, single-cell clones were sorted into 96-well cell culture plates using the BD FACSAria^TM^ Fusion Cell Sorter (BD Biosciences), before screening for the loss of ANXA1 expression using western blot analysis (**Extended Data Fig. 9a**).

ANXA2 CRISPR/Cas9 KO (sc-419418) and homology-directed repair (sc-419418-HDR) plasmid constructs were obtained from Santa Cruz Biotechnology (SCBT), and co-transfected into 4T1 cells using Lipofectamine^TM^ 3000 reagent (Invitrogen). 48 hours post transfection, cells were selected using puromycin at 5 µg/mL for 3 days. Subsequently, single-cell clones were sorted into 96-well cell culture plates using the BD FACSAria^TM^ Fusion Cell Sorter (BD Biosciences), before screening for the loss of ANXA2 expression using western blot analysis (**Extended Data Fig. 9b**).

### Western blotting

Western blots were performed to verify the protein expression level of different breast cancer cell lines. For each cell line, 3 × 10^5^ cells were seeded onto 6-well cell-culture treated plates and incubated for 24 hrs at 37°C in a 5% CO_2_ humidified incubator. Seeded cells were harvested in RIPA buffer (Pierce Biotechnology) with protease/phosphatase inhibitor cocktail (Pierce Biotechnology). Cell lysates were clarified by centrifugation at 17,000 x g for 20 min at 4°C. Protein concentrations of the samples were quantified against reconstituted bovine serum albumin (BSA) (Pierce Biotechnology) of known concentrations using Bradford reagent (Bio-Rad). Samples (15 µg) were separated on a 4 – 15% PAGE gel (Bio-Rad) and transferred onto 0.2 µM nitrocellulose membrane (Bio-Rad). Membranes were blocked with 5% BSA (VWR) phosphate-buffered saline with 0.1% Tween-20 (PBST) for 1 hr at room temperature before incubating with primary antibodies specific for ANXA1 (1:1000; ab65844, Abcam), ANXA2 (1:1000; sc-28385, SCBT), β-actin (1:2000; A1978, Sigma-Aldrich), and β-tubulin (1:2000;#2146, Cell signaling technology) as well as TRPM7 (1:1000; ab262909, Abcam) and β-actin (1:2000; A1978, Sigma-Aldrich) overnight at 4°C. Subsequently, membranes were incubated in secondary horseradish-peroxidase-conjugated mouse-specific (1:5000; P0260, DAKO) and rabbit-specific antibodies (1:5000; PI-1000, DAKO) for 1 hr at room temperature. Bands were visualised with Clarity^TM^ Western ECL Substrate (Bio-Rad) imaged using ChemiDoc^TM^ MP Imaging System (Bio-Rad).

### Visualization of actin structure and analysis of actin anisotropy

To dynamically track cell actin structure, we created MDA-MB-231 and MCF10A cell lines stably expressing LifeAct-TdTomato. The LifeAct-TdTomato^56^ (a gift from L. Rong) plasmids were purchased from Addgene and used for lentiviral cell transduction. Fluorescence activated cell sorting (FACS) were performed to sort for upper 20% of cells with higher fluorescent signal of LifeAct-TdTomato. The sorted cells were then cultured following the standard cell culture process. For actin visualization, cells were introduced into microfluidic devices following the process of microfluidic assay. Subsequently, the microfluidic devices were transferred to the W1 confocal microscope equipped with a live-cell incubator system. To minimize the potential impact of phototoxicity, imaging was conducted at -1 min (before flow introduction), 1 min (after flow introduction), and every 10 mins thereafter until the experiment concluded.

For analyzing actin dynamic, the ImageJ-based FibriTool, as described in previous work^57^, has been employed. This plug-in estimate actin anisotropy based on user-defined areas and line segments. The anisotropy measurements were obtained using the FibriTool plug-in

### Intravital experiments in mouse

An illustration for briefly describing has been shown in **Extended Data Fig. 10a**. For tumor cell imaging, *in vitro* cultured MDA-MB-231 cells were removed culture medium and washed with PBS. Following the commercial protocol (CellTracker^TM^ Deep Red, C34565, Thermos Fisher), PBS mixed with 0.5 μg/ml were cultured with adhesive cells for staining at 37°C cell culture incubator, 15-20 min. The mixed PBS solution was removed and cells were washed by fresh PBS. The subsequent process followed the cell culture method until receiving cell pellets. Membrane-labeled MDA-MB-231 cells (∼2 ×10^6^) were resuspended in 200 μL PBS. Then, the C57BL/J mice (male, 4∼8 weeks) were anesthetized and dissected to expose the spleen. PBS with cells injected into the mice through spleen injection. The wound was mended. Then, 70kDa dextran (Texas Red, D1830, Thermos Fisher, 1mg / mouse) was injected into mice by intravenous injection. After 1 hour, the mice were re-anesthetized and dissected to expose the liver on a plate with a #1.5, 1.5, F12 mm, round cover glass in the center for intravital imaging. During imaging, the mice body was fixed in an artificial device and its body temperature was maintained around ∼37 °C.

All experimental procedures were approved by the Institutional Animal Care and Use Committee at Tsinghua University, Beijing, China.

### Viability analysis

Caspase 3 (NucView® 488 Caspase 3 Assay Kit, 30029-T, Biotium) was used to determine cell viability. Cell death was deemed once cell express caspase 3 signal and become totally rupture appear within few frames (**Fig. 6a and 6b**). All the viability results were measured based on at least 3 independent chip experiments with hundreds of cells.

### Statistical analysis

Data are presented as the mean ± s.d. from ≥ 3 independent experiments (microfluidic chips), representing different biological replicates, with ≥ 20 cells per condition per experiment (≥ 10 cells for actin anisotropy). Data points indicate values from individual cells, unless stated otherwise. Shapiro-Wilk tests were conducted for data expressed as percentages where the number of data points ranged from 3 to 6. Datasets following a Gaussian distribution were compared using a two-tailed Student’s t-test or one-way ANOVA followed by Tukey’s post hoc test. *P* > 0.05 was considered not statistically significant (n.s.); *P* < 0.05 was considered statistically significant, with \**P* < 0.05, \*\**P* < 0.01, and \*\*\**P* < 0.001.

### Figure materials

Part cartoon images are obtained from BioRender with available license.

## Acknowledgements

This work was supported by the Institute for Health Innovation and Technology (iHealthtech) (No. A-0001415-06-00), and the Mechanobiology Institute (No. A-0003467-20-00) at the National University of Singapore (NUS). We appreciate the support from the NUS Startup Grant (No. A-8001301-00-00) provided to MechanoBioEngineering Laboratory at the Department of Biomedical Engineering at NUS.

## Author contributions

L.L. conceptualized and designed this study, performed all *in vitro* microfluidics-based experiments and analysed data, selected *in vivo* experiments and analysed *in vivo* date, as well as writed the manuscript. Z.Z. and Y. C. performed most *in vivo* experiments including mouse anatomy and intravital imaging, as well as analysed *in vivo* date. Jonathan C. responded to the generation of Annexin-associated knockout cell line and western blot analysis. X.S. performed selected microfluidics-based experiments and flow cytometry-associated experiments, analysed data, as well as edited the manuscript. Lina L. supervised the Annexin-associated knockout experiemnt and edited the manuscript. J.W. and Q.D. supervised the intravital imaging and edited the manuscript. A. H. and C.L. conceptualized, designed and critically supervised this study, and wrote the manuscript

---

**Extended Data Fig. 1.**
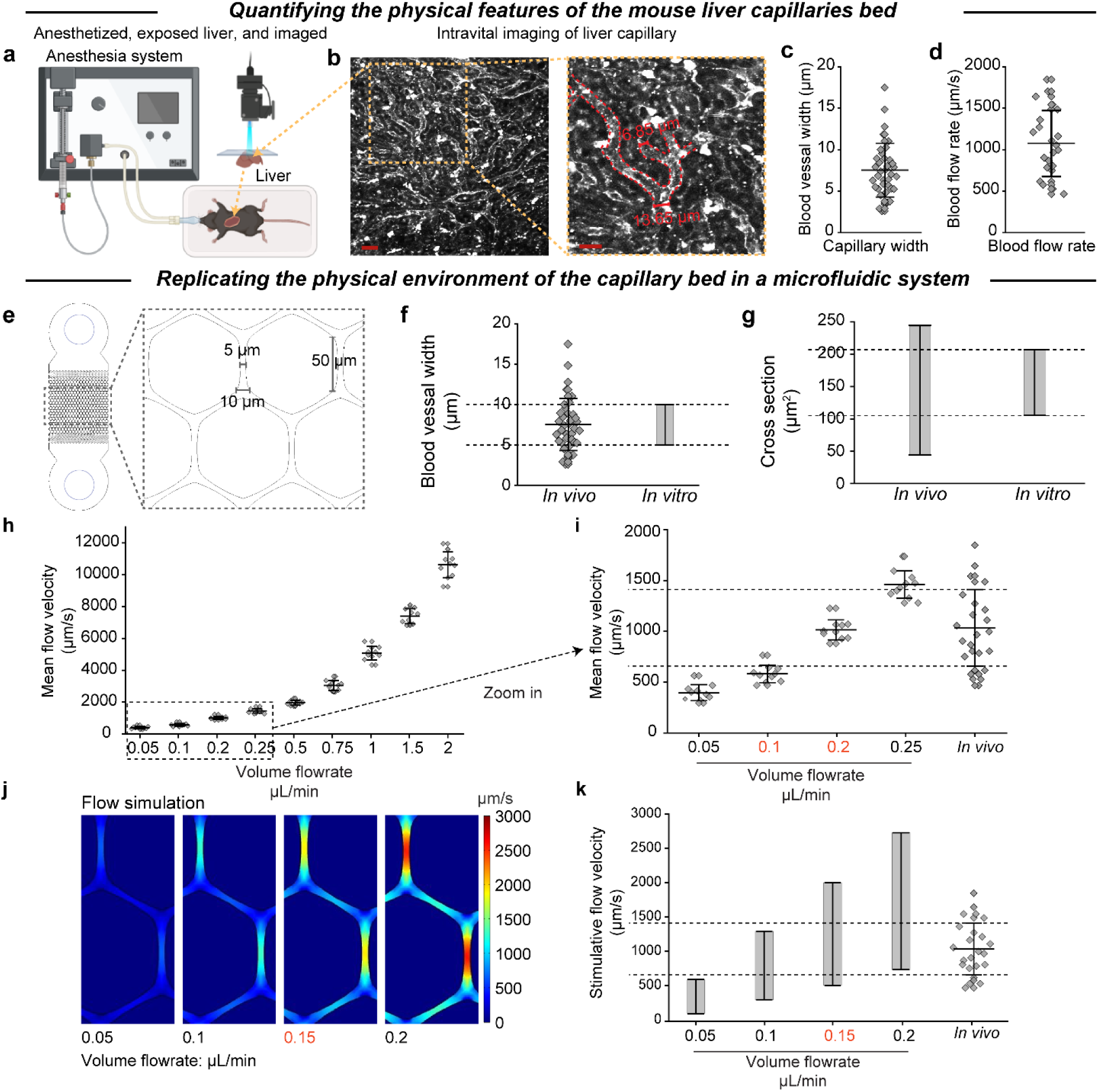
From *in vivo* to *in vitro*: replicating capillary bed in a ‘capillary bed-on-a-chip’ microfluidic device. **a-b**, *In situ* intravital imaging of mouse liver capillary vessels. Scale bars, 20 μm. **c-d**, Quantification of physical characteristics, including diameter (**c**) and flow rate (**d**), in blood vessels. **e-k,** Schematic representation of ‘capillary bed-on-a-chip’ microfluidic device. This system replicates *in vivo* physiological characteristics, including blood vessel width (**f**), cross-sectional shape (**g**), alongside their *in vitro* topological parameters, and flow rate (**h-k**). **h and i**, a microbeads-based method to visualize flow velocity within the chip. **j and k**, a COMSOL-based method to measure the flow velocity distribution throughout the chip. Data are represented as the mean ± standard deviation (n ≥ 3 independent chip experiments).

**Extended Data Fig. 2.**
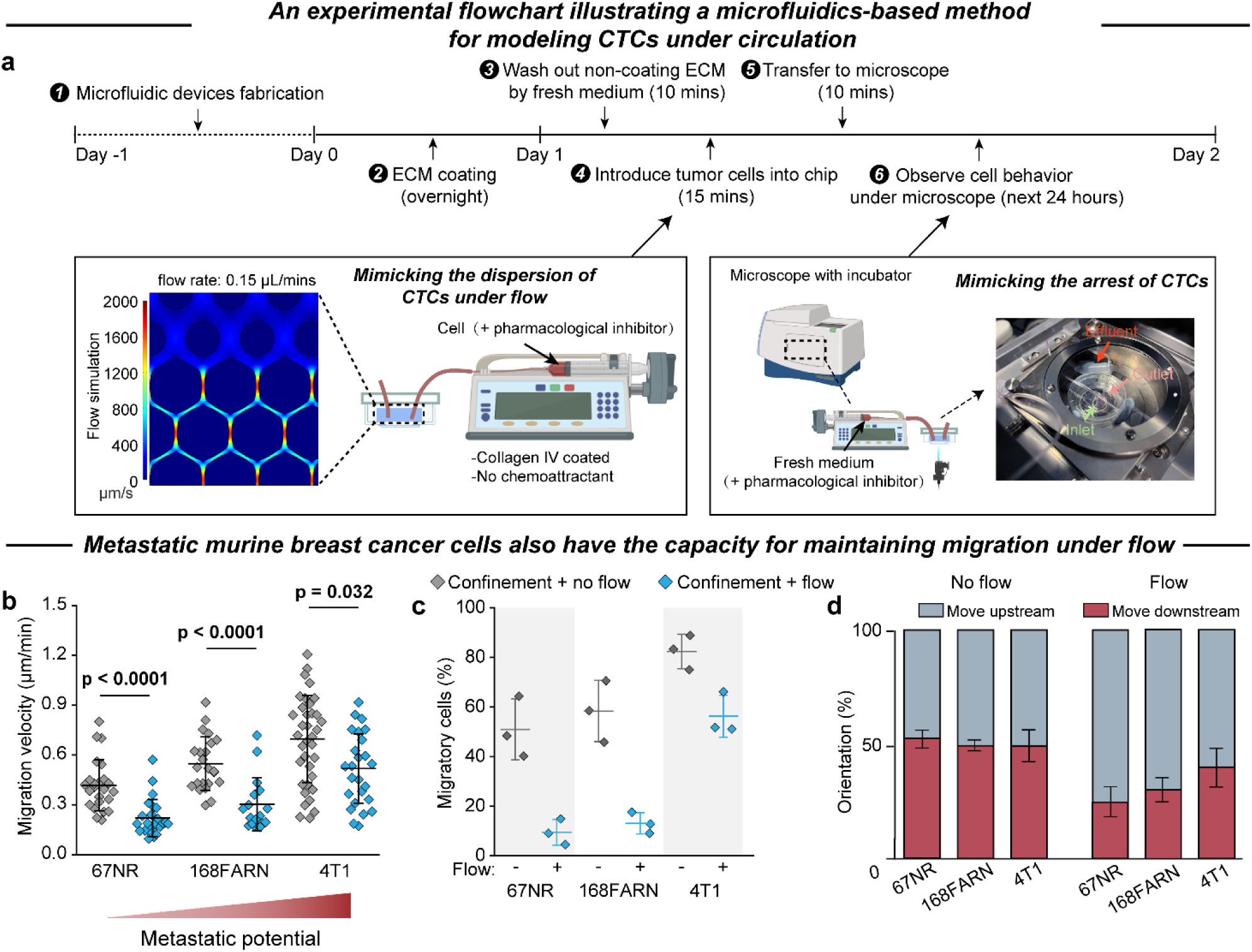
Schematic of microfluidic experiments to study cell migration. **a**, Experimental timeline of the ‘capillary bed-on-a-chip’ microfluidic system to study tumor cell migration. **b-d**, Analysis of the flow-insensitive migration on murine mammary gland cells. Analysis of the migration speed (**b**), migratory cells (**c**), and migration orientation preference (**d**) on murine mammary gland tumor cells. Data are represented as the mean ± standard deviation (n ≥ 3 independent chip experiments). Statistical analysis was performed using one-way ANOVA followed by Tukey’s test (**b**).

**Extended Data Fig. 3.**
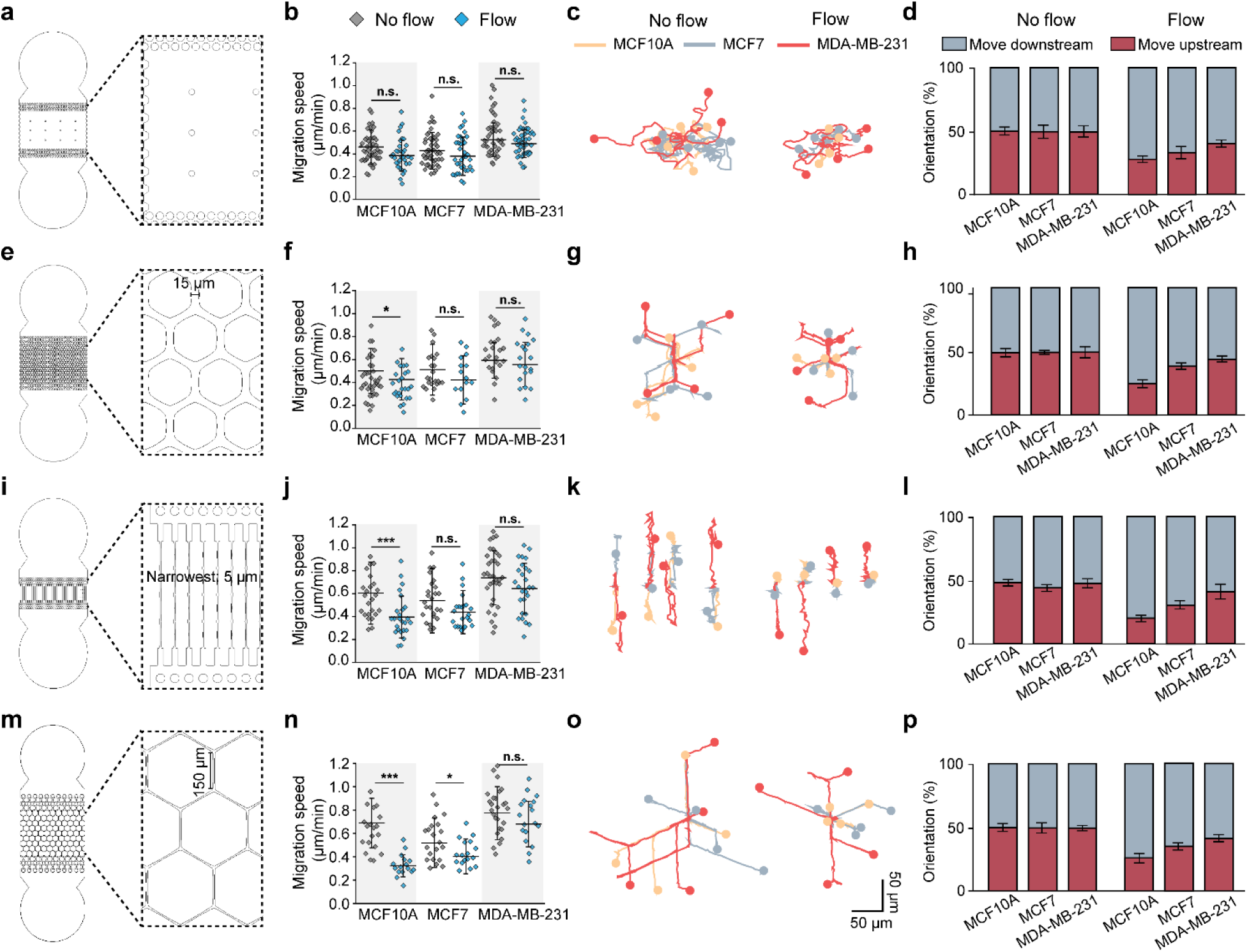
Flow-insensitive migration, associated with metastatic potential, is a common phenomenon under flow and confinement condition. **a-d**, Analysis of breast cancer cell migration within a non-confined microfluidic chip. **e-h**, Analysis of breast cancer cell migration within a minimally confined microfluidic chip. **i-l**, Analysis of breast cancer cell migration within a line-confined microfluidic chip. **m-p**, Analysis of breast cancer cell migration within a capillary-like microfluidic chip featuring an extended channel length. Data are represented as the mean ± standard deviation (n ≥ 3 independent chip experiments). Statistical analysis was performed using one-way ANOVA followed by Tukey’s test (**b, f, j** and **n**). P values for comparisons with the results for intracellular calcium dynamic between no flow and other two conditions are indicated (\**P* < 0.05; \*\*\**P* < 0.001). N. S., not significant. Scale bars, 50 μm.

**Extended Data Fig. 4.**
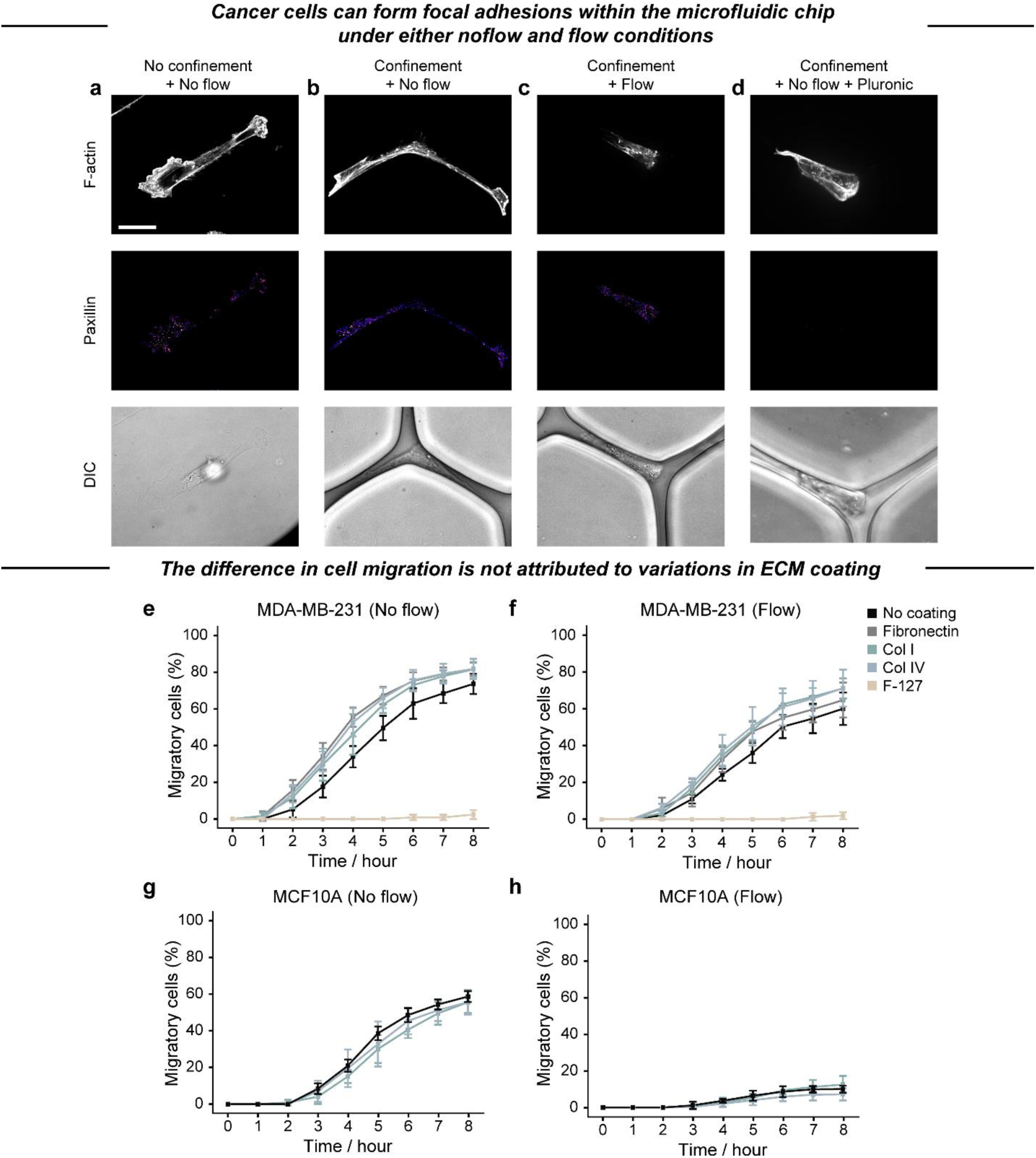
Formation of focal adhesion and the effect of ECM proteins on flow-insensitive migration. **a-d**, Confocal imaging of actin structures and paxillin-based focal adhesions in MDA-MB-231 cells cultured on a glass-bottom dish (**a**), within a microfluidic chip without flow (**b**), with flow (**c**), and in a chip treated with an anti-focal adhesion agent pluronic F-127 (**d**). Scale bars, 20 μm. **e and f**, Time-dependent analysis of MDA-MB-231 cell migration in the presence of various ECM proteins. **g and h**, Time-dependent analysis of MCF10A cell migration in the presence of various ECM proteins. Data are represented as the mean ± standard deviation (n ≥ 3 independent chip experiments).

**Extended Data Fig. 5.**
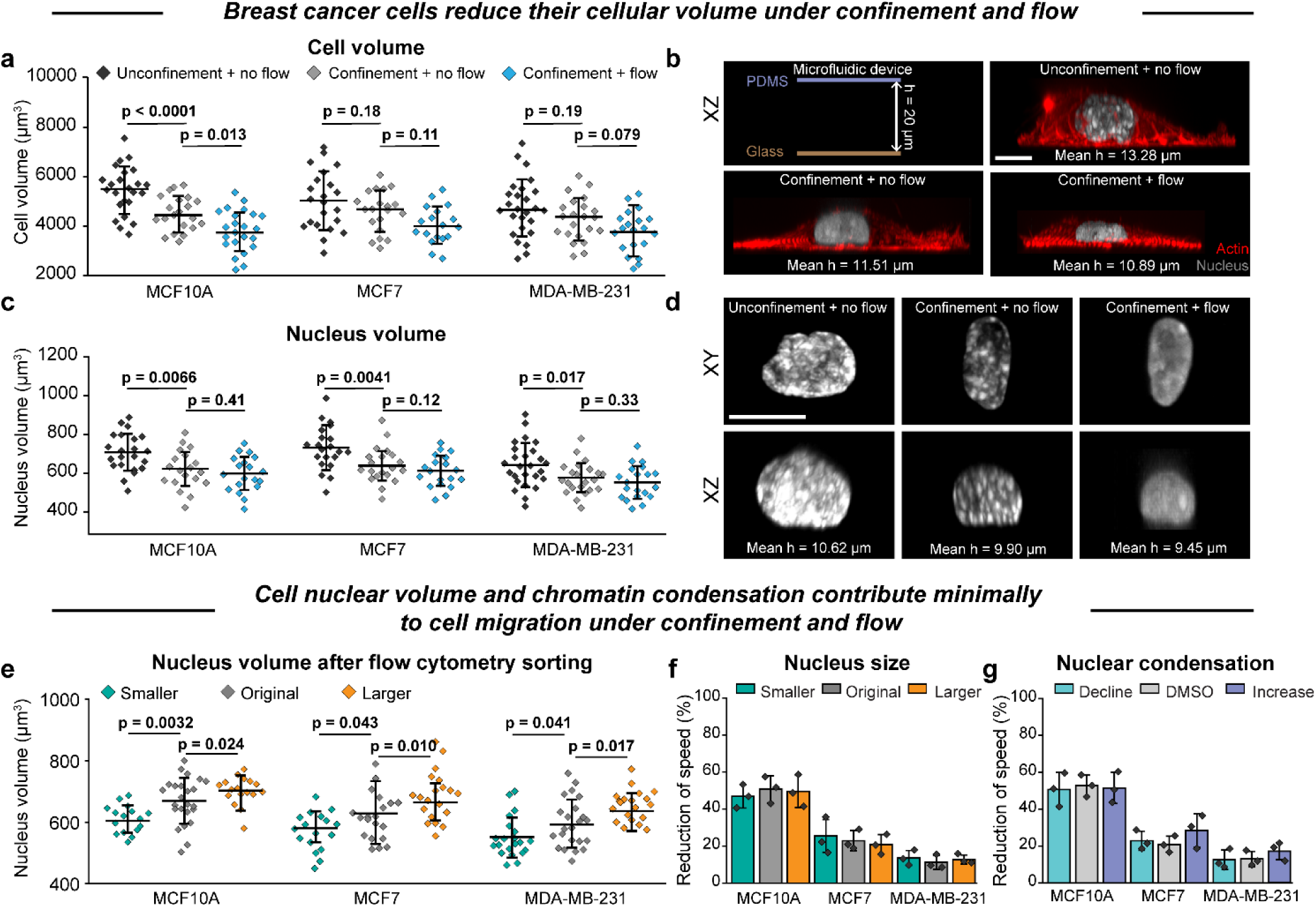
Effect of nucleus difference on cell migration under confinement and flow. **a and b**, Analysis of cell volume and cell height via confocal imaging. **c and d**, Analysis of nucleus volume and nucleus height via confocal imaging. **e**, Assessment of nucleus volume following flow cytometry sorting. **f**, Reduction in migration speed under flow of breast cancer cell with varying nuclear sizes. **g**, Reduction of speed under flow is not related with either reducing chromatin condensation (RGFP-966) or increasing chromatin condensation (JIB-04). Data are represented as the mean ± standard deviation (n ≥ 3 independent chip experiments). Statistical analysis was performed using one-way ANOVA followed by Tukey’s test (**a, c** and **e**). All scale bars, 20 μm.

**Extended Data Fig. 6.**
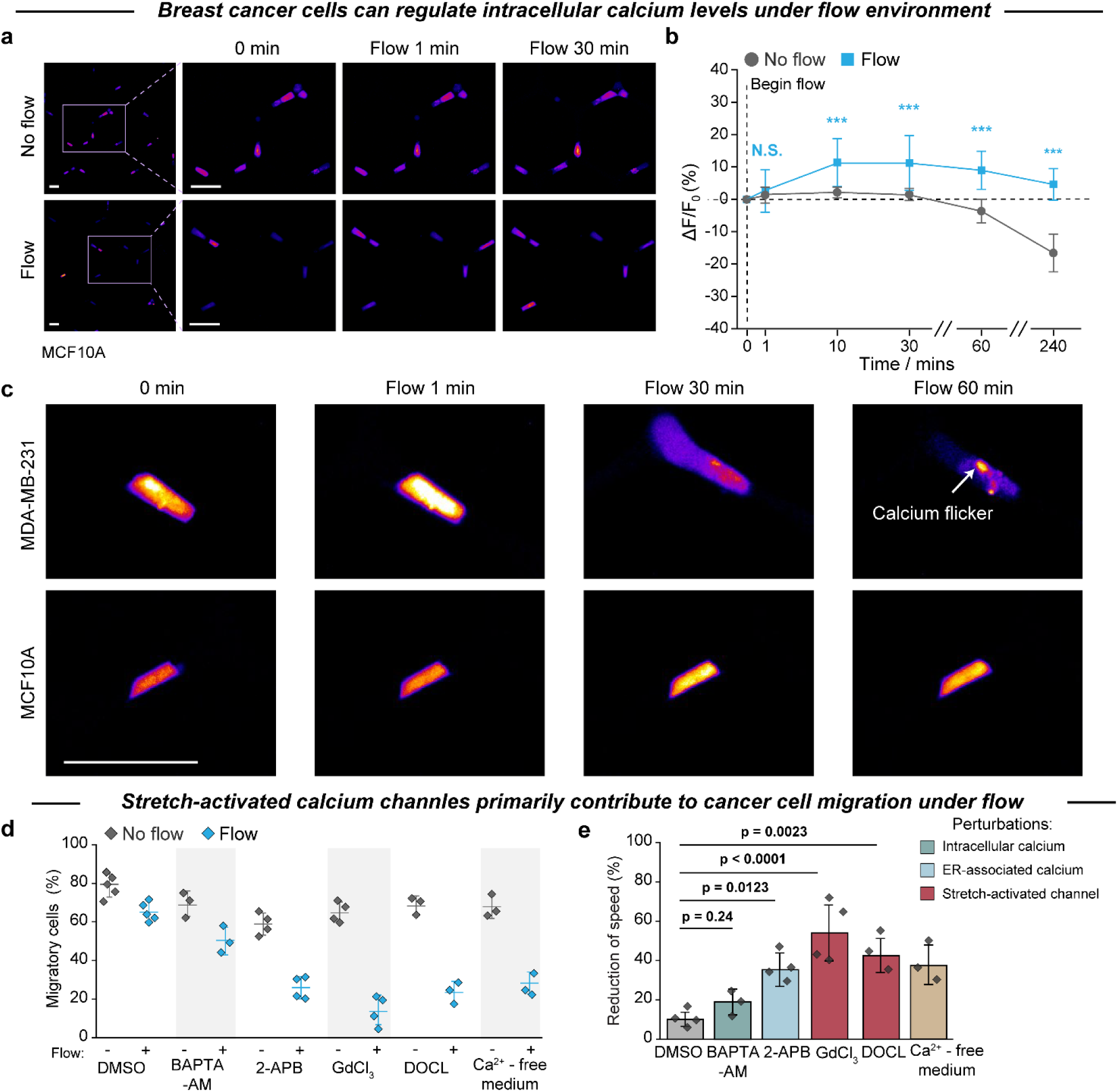
The roles of calcium in cell mechanotransduction in response to flow. **a and b**, Quantification of dynamic changes in intracellular calcium in MCF10A cells. **c**, Calcium flickers typically occur on MDA-MB-231 cells but rarely on MCF10A cells. **d and e,** Disruption of calcium-related pathways using pharmacological inhibitors and their effects on cell motility. For detailed information on each inhibitor, refer to the Methods section under pharmacological treatments. Data are represented as the mean ± standard deviation (n ≥ 3 independent chip experiments). Statistical analysis was performed using Kruskal-Wallis tests followed by Dunn’s test (**b** and **e**). \*\*\**P* < 0.001, N. S., not significant. All scale bars, 50 μm.

**Extended Data Fig. 7.**
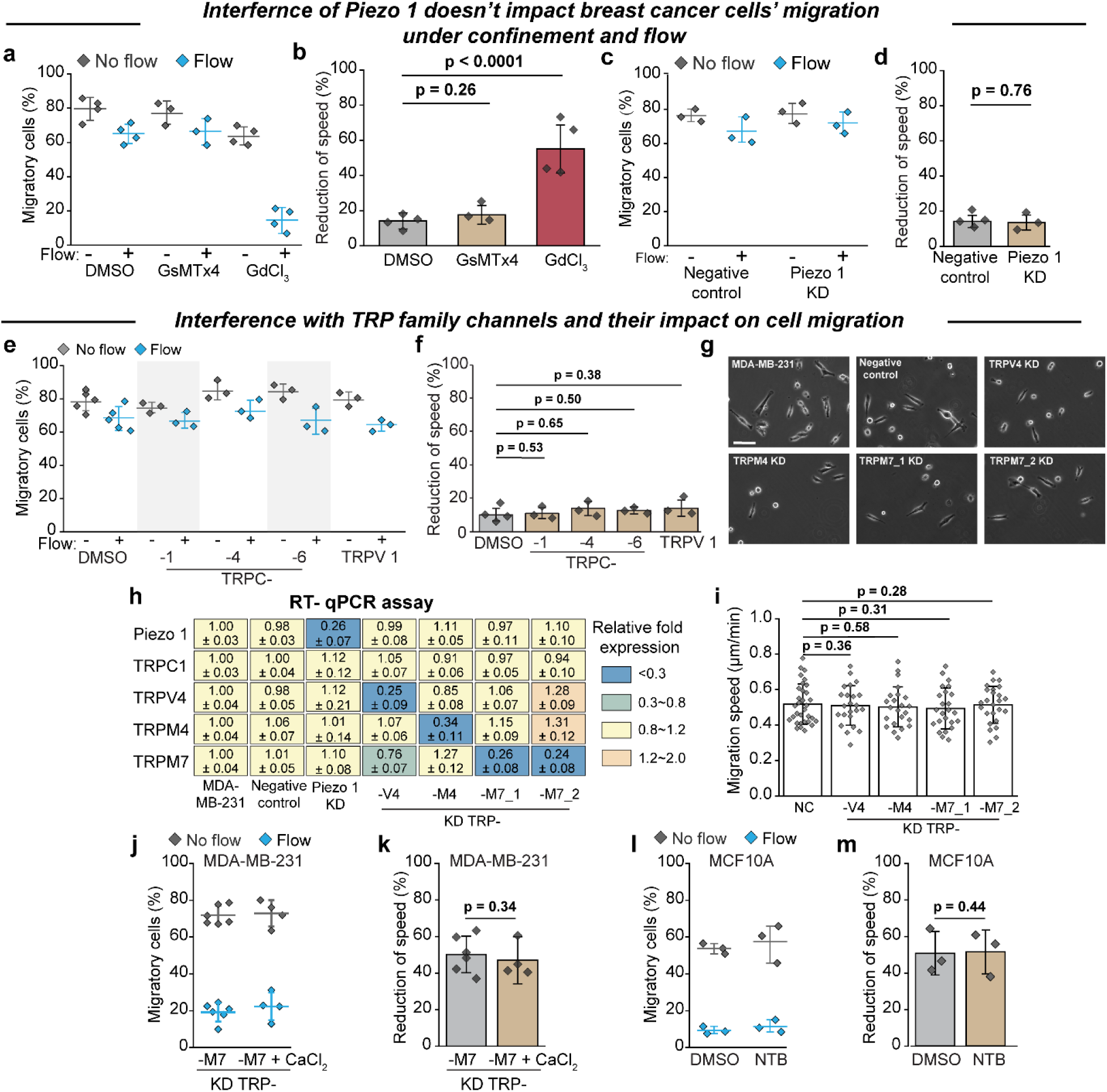
The roles of calcium-related ion channels in cell migration under confinement and flow. **a-d**, The roles of Piezo 1 channel in cell migration. **a and b**, A peptide-based method (GsMTx4) to inhibit the function of Piezo 1 and its impact of cancer cell motility. **c and d,** Analysis of the motility on MDA-MB-231 cancer with Piezo 1 knockdown. e-m, The roles of TRP family channels in cell migration. **e and f**, Pharmacological inhibition of TRP family proteins and its effect on MDA-MB-231 cell migration. **g**, Bright-field imaging to compare MDA-MB-231 cell morphology following siRNA knockdown. Scale bars, 20 μm. **h,** Quantification of gene expression following various siRNA knockdowns using RT-qPCR. **i,** Measurement of MDA-MB-231 cell migration speed after different siRNA knockdowns. **j and k,** Increasing extracellular calcium concentration does not impact migration in MDA-MB-231 cells with TRPM7 knockdown. **l and m**, Activation of the TRPM7 channel using naltriben (NTB) does not change the flow-induced non-migratory state of MCF10A cells. Data are represented as the mean ± standard deviation (n ≥ 3 independent chip experiments and 3 independent gene extraction). Statistical analysis was performed using Kruskal-Wallis tests followed by Dunn’s test (**b, d, f, k** and **m**) and one-way ANOVA followed by Tukey’s test (**i**).

**Extended Data Fig. 8.**
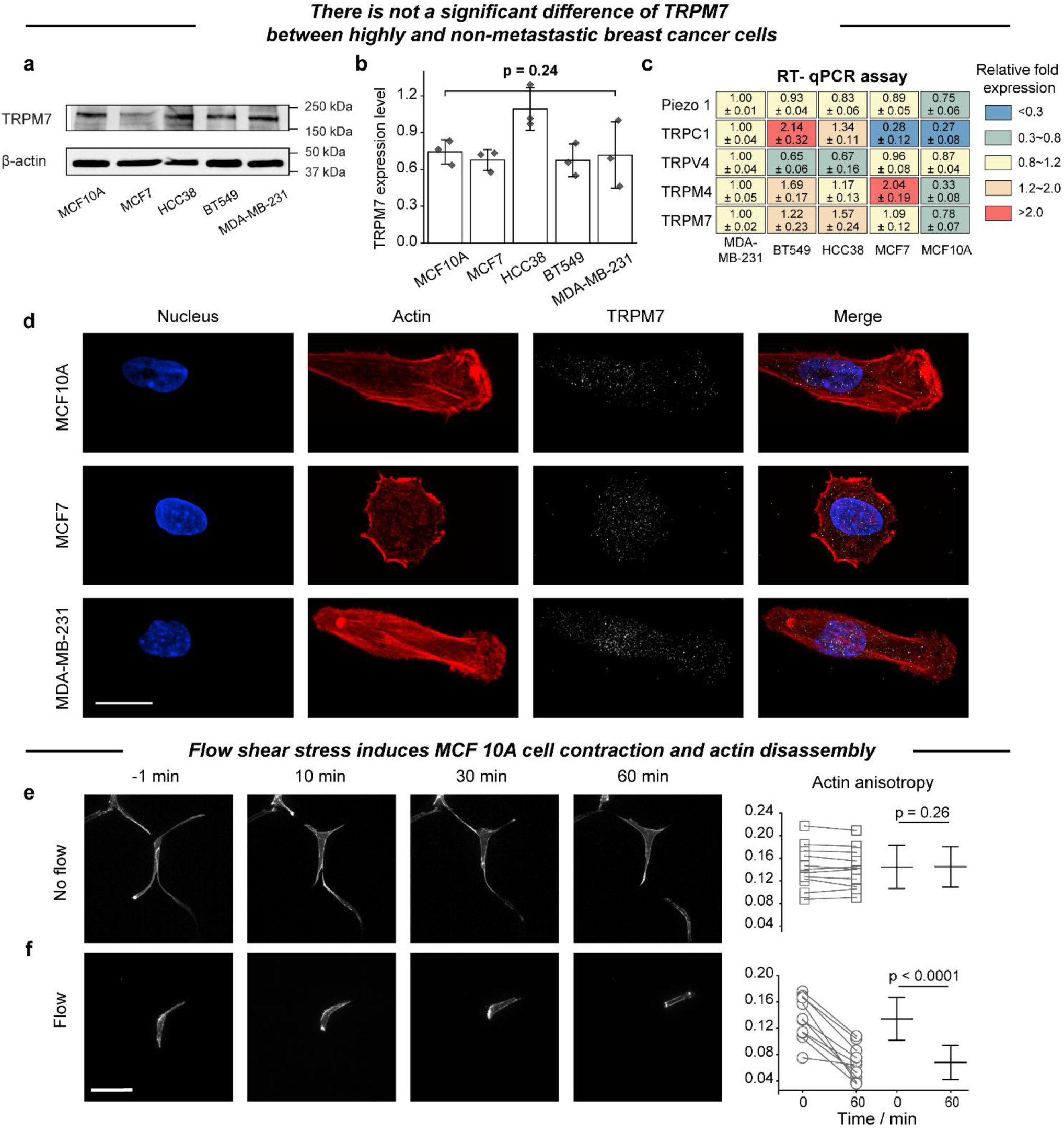
Analysis of TRPM7 in metastatic vs. non-metastatic breast cancer cells. **a and b**, Quantification of TRPM7 expression level across a panel of breast cancer cell lines via western blot. **c**, Assessment of the expression levels of various ion channels using RT-qPCR. **d**, Immunofluorescence analysis to examine the expression and distribution of TRPM7. Scale bars, 20 μm. **e-f**, Confocal imaging to analyze actin dynamic in MCF10A cell. Scale bars, 50 μm. Data are represented as the mean ± standard deviation (3 independent gene or protein extraction). Statistical analysis was performed using Kruskal-Wallis tests followed by Dunn’s test (**b**) and one-way ANOVA followed by Tukey’s test (analysis of actin anisotropy in **e** and **f**)

**Extended Data Fig. 9.**
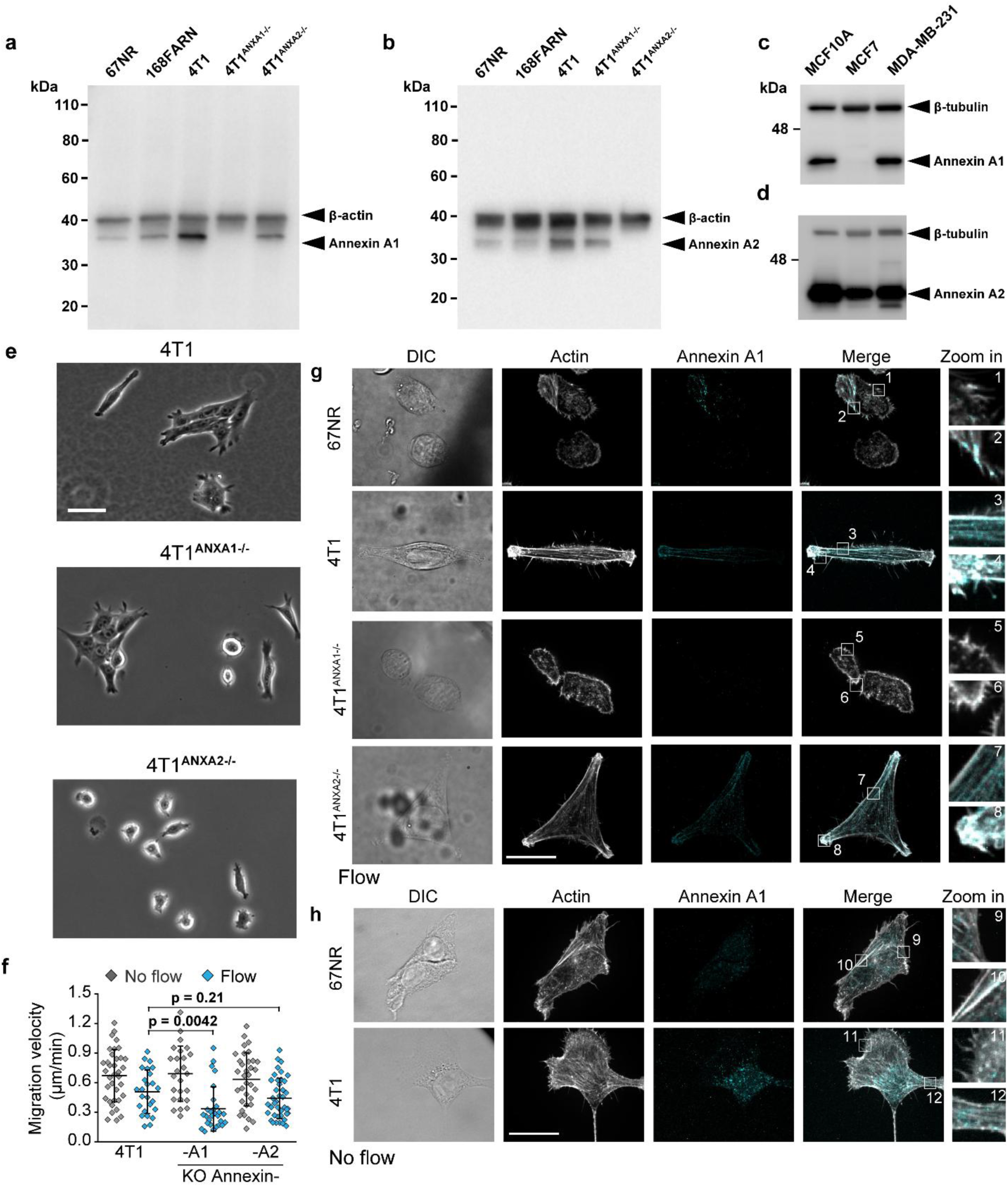
The effect of Annexin A1 and A2 knockout on cell migration under flow condition. **a-d**, Original blots to verify the expression of Annexin A1 and A2 for breast cancer cells with different metastasis potential. **e**, Bright-field imaging to compare the morphology of 4T1 cells and their Annexin knockout subpopulation. Scale bars, 50 μm. **f**, Analysis of the migration velocity on two 4T1 cells Annexin knockout subpopulation. **g and h**, Immunofluorescence analysis to examine the co-localization of actin and Annexin A1 under flow (**g**) and no flow (**h**). Scale bars, 20 μm. Statistical analysis was performed using one-way ANOVA followed by Tukey’s test (**f**).

**Extended Data Fig. 10.**
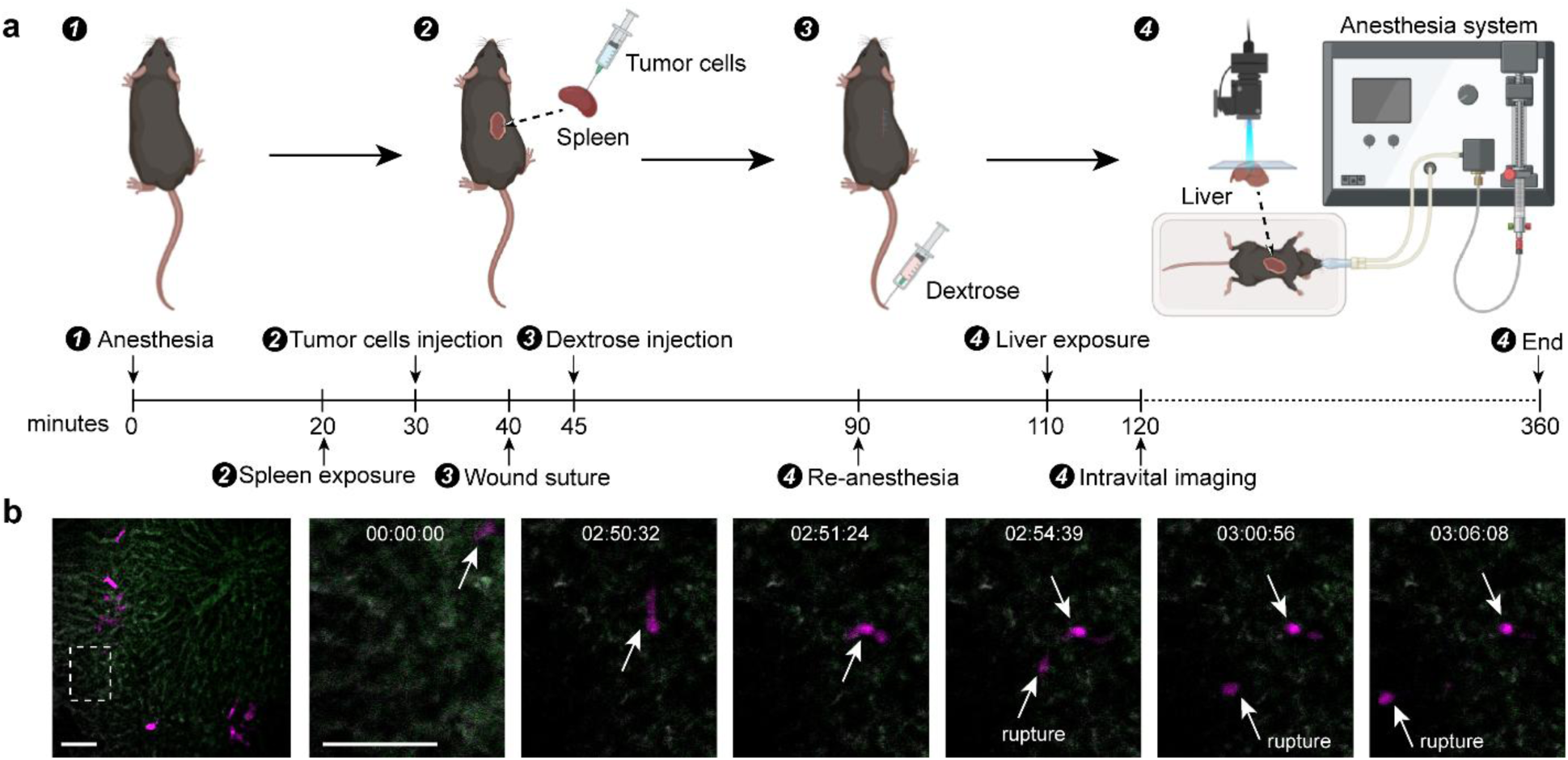
Intravital imaging of MDA-MB-231 cells within mouse liver capillary vessels. **a**, Schematic representation of the intravital imaging process. **b,** Observation of cancer cell rupture within liver capillary. Scale bars, 50 μm.

**Extended Data Fig. 11.**
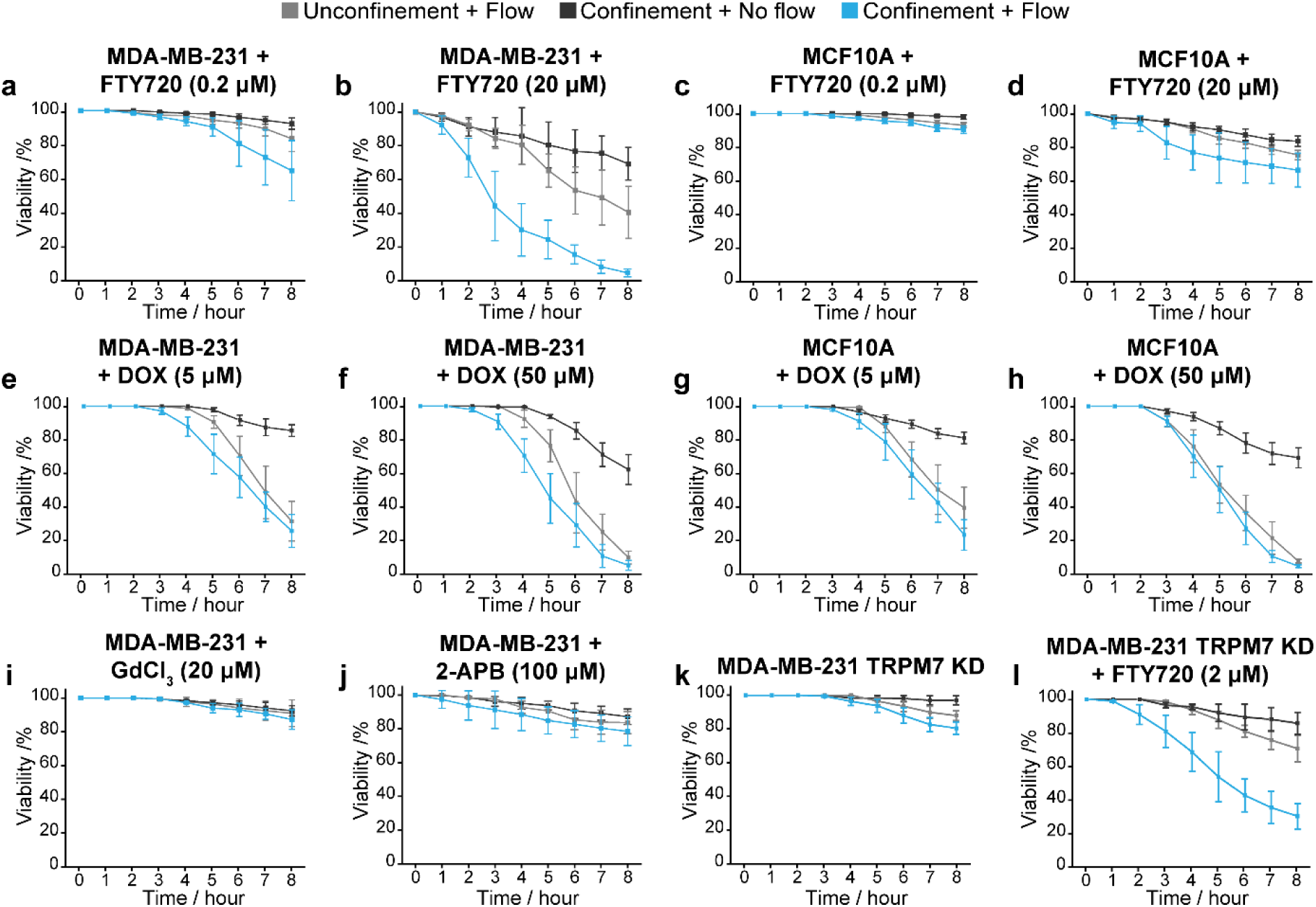
Cell viability under drug treatment and physical stresses. **a-b**, Cell viability of metastatic cancer cells MDA-MB-231 treated with concentration gradient of FTY720. **c-d**, Cell viability of non-tumorigenic cells MCF10A treated with concentration gradient of FTY720. **e-h**, Traditional chemotherapy drug doxorubicin is highly poisonous for both cancer cells and non-tumorigenic cells. Positive control. **i-j**, Cell viability of metastatic cancer cells MDA-MB-231 treated with calcium-associated inhibitor. **k-l**, Cell viability of metastatic cancer cells MDA-MB-231 with TRPM7 knockdown treated with concentration gradient of FTY720. All data are represented as the mean ± standard deviation (n ≥ 3 independent chip experiments).

